# Electrical stimulation of the superior temporal gyrus evokes rapid responses in human visual cortex

**DOI:** 10.1101/2025.03.22.644725

**Authors:** Emily C. Cunningham, David Brang

**Affiliations:** Department of Psychology University of Michigan 530 Church St. Ann Arbor, MI, 48109

## Abstract

Sounds elicit rapid responses in human visual cortex. Anatomical work in nonhuman primates suggests that these responses may be enabled by monosynaptic, corticocortical projections between auditory and visual areas. Such projections would not only provide routes for rapid modulation of visual processing in the sighted, but also avenues for cortical adaptation following vision loss. However, there is little available information on the presence and organization of such projections in humans. Here, we address this question by examining intracranial responses to direct electrical stimulation of the superior temporal gyrus (STG) in 23 patients (both male and female). Mid- and posterior-STG stimulation produced rapid responses in early visual cortex (18ms+), with a distribution that favored lateral occipital cortex and anterior calcarine regions near peripheral V1. Early visual cortex (V1/V2) responded most strongly to stimulation over mid-STG, whereas lateral occipital cortex (near V5/hMT+) exhibited the most robust response to posterior STG stimulation. These data demonstrate that communication from auditory to visual cortex can occur in humans at latencies that are compatible with corticocortical, potentially monosynaptic, transmission. Responses in visual cortex were organized in a manner similar to that of nonhuman primates, with preliminary evidence for an exception over the occipital pole. Overall, the results provide support for theories that human visual responses to sound are inherited, in part, from auditory cortex.

## Introduction

Early visual cortex is increasingly recognized as a site of rapid audiovisual interaction. Even in the absence of concurrent visual stimulation, sounds modulate activity in human visual cortex within 100ms of onset (Brang et al., 2015; Ferraro et al., 2020; Mercier et al., 2013; Plass et al., 2019; Raij et al., 2010), with responses as early as 30ms recorded in the anterior portion of human V1 (Brang et al., 2015). These responses are thought to indicate a sound-related alignment of visual cortical activity that enhances sensitivity to subsequent visual input (Brang et al., 2015; Lakatos et al., 2009; Mercier et al., 2013; Naue et al., 2011; Plass et al., 2019; Romei et al., 2012; see also Thorne & Debener, 2014). However, relatively little is known regarding the pathways by which these signals are communicated in humans. This information is not only necessary for a complete understanding of how sounds modulate the representation of visual information (e.g. Frassinetti et al., 2002; McDonald et al., 2000; Noesselt et al., 2010; Shams et al., 2000; Vroomen & Gelder, 2000), but also augments our understanding of the routes by which visual cortex can be recruited by sounds following visual deprivation. Nonhuman primate tracing studies point to several potential avenues for cross-sensory communication, including subcortical routes that bypass auditory cortex (possibly through the tectopulvinar system; Lyon et al., 2010), corticocortical pathways between early sensory areas (Falchier et al., 2002; Rockland & Ojima, 2003), and/or polysynaptic routes from auditory cortex through multisensory association areas or thalamic intermediaries (Borra & Rockland, 2011; Cappe et al., 2009; Clavagnier et al., 2004). In addition, sounds may exert indirect effects on visual cortical activity by initiating processes associated with arousal, attentional capture, and eye movements. Of these candidates (all of which may be present to some extent in humans), particular recent interest surrounds the possibility that human visual cortex may receive direct (i.e., corticocortical) input from primary or early auditory association cortex.

Some indirect support for the existence of such projections in humans can be gleaned from the features of sound-evoked visual responses. First, the onset of the response in visual cortex only slightly exceeds that of the earliest recorded responses in auditory cortex (by between 2-15ms; Brang et al., 2015; Raij et al., 2010). This speed suggests relatively direct communication, consistent with either a subcortical or early corticocortical relay. It is difficult to differentiate between these two routes (corticocortical vs. subcortical) by response profile alone, and contributions of these pathways are not mutually exclusive. However, there is some additional evidence favoring cortical over subcortical origins. For example, whereas subcortical auditory neurons typically entrain to amplitude-modulated (AM) stimuli, a large portion (∼20%) of neurons in auditory cortex respond only to onset/offset of an AM stimulus train (Bieser & Müller-Preuss, 1996; see also Harms & Melcher, 2002). Consistent with what one would expect of responses inherited from an auditory cortical source, visual cortex is sensitive to sound onset/offset, but does not passively synchronize or entrain to auditory amplitude modulation (Brang et al., 2022; see also Cunningham et al., 2022). Taken together with the speed of the initial sound-evoked visual response, this profile suggests communication from early auditory cortex.

In addition, there are suggestive similarities between the distribution of human sound-evoked visual responses and patterns of nonhuman primate connectivity. For example, in monkeys, monosynaptic projections from auditory to early visual cortex are biased toward the visual periphery (Falchier et al., 2002; Rockland & Ojima, 2002), with some projections also identified between superior temporal and visual motion sensitive areas (Majka et al., 2019; Palmer & Rosa, 2006). In humans, occipital responses to sound emerge broadly across visual cortex, with some concentration in lateral occipital regions (near human V5/hMT+; Mercier 2013; Plass 2019), and notably fast responses in anterior portions of the calcarine sulcus (Brang et al., 2015; Ferraro et al., 2020).

Although the distribution, specificity to onset/offset, and speed of sound-evoked visual responses are consistent with communication from auditory cortex, these observations are at best indirect. Some progress in linking nonhuman primate anatomical observations with human sound-evoked visual responses has been made through noninvasive diffusion-based imaging (DTI), with at least one report identifying possible connections between primary auditory cortex and both the anterior calcarine sulcus and occipital pole (Beer et al., 2011), and another detailing connections between planum temporale and visual motion-sensitive hMT+ (Gurtubay-Antolin et al., 2021). However, limits to the sensitivity of DTI and absence of directional/causal information restrict the conclusions that can be drawn from this approach. Given the sparse nature of the projections identified in macaques, the question of whether (and how) projections from auditory to visual cortex manifest in humans remains difficult to resolve.

Here, our aim is to establish and describe the pattern of directional communication from human auditory to visual cortex. One way to approach this question is to examine the responses evoked in human visual cortex following direct electrical stimulation of regions in and around auditory cortex. To that end, we take advantage of intracranial EEG (iEEG) recordings made during administration of single-pulse electrical stimulation (SPES) in patients with epilepsy (open data from van Blooijs et al., 2023). SPES provides an opportunity to estimate effective connectivity between brain regions through the distribution/latency of responses evoked when electrical current is passed between adjacent intracranial contacts. When stimulation and recording sites are located on the cortex, such responses are termed corticocortical evoked potentials, or CCEPs (Matsumoto et al. 2004; for reviews of the technique, see Keller et al., 2014; Matsumoto et al., 2017). CCEP waveforms vary in shape, but often display a rapid initial deflection (usually <50ms, though longer latencies have been reported, e.g. Araki et al., 2015) along with a subsequent slower wave (∼80-250ms; Figure 1A). The first-emerging peak in this signal (typically referred to as ‘N1’, though it may be positive or negative) has been used to characterize transmission speed based on the assumption that it results from orthodromic communication along corticocortical pathways between stimulation and recording sites (e.g. Lemaréchal et al., 2022; van Blooijs et al., 2023). This assumption derives largely from reports that the latency of this peak scales linearly with surface distance from the stimulation site (e.g. (Matsumoto et al., 2012; Trebaul et al., 2018; Yamao et al., 2014), together with comparisons of CCEPs to responses following stimulation of white matter fiber bundles (Yamao et al., 2014). Although other features of the CCEP waveform can provide additional information regarding interregional communication (e.g. Huang et al., 2023; Miller et al., 2021), here we focus on the presence and latency of the initial ‘N1’ peak, building from the assumption that its distribution and timing provide insight into the distribution of directional projections from auditory to visual cortex.

**Figure 1.**
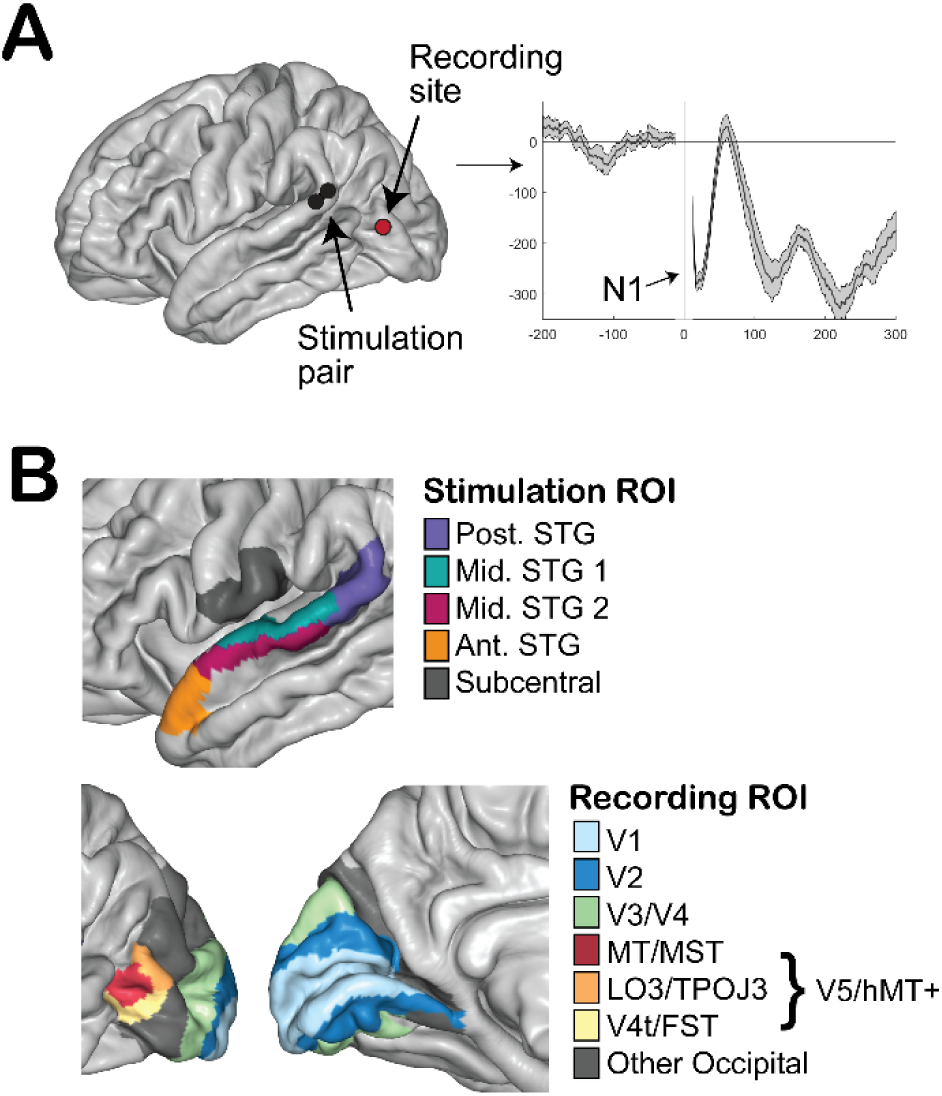
(A) Representative CCEP response with prominent early negative deflection (N1) elicited by stimulation of the posterior STG, and (B) regions of interest in the superior temporal gyrus, subcentral gyrus (control), and occipital lobe illustrated on the Freesurfer average brain.

The CCEP approach provides a test of the question of whether (and to what extent) rapid communication from auditory to visual cortex occurs in humans. Evidence consistent with corticocortical communication (in the form of low-latency occipital responses following stimulation of auditory cortex) would support theories that the initial visual response to sound is relayed, at least in part, from auditory cortex (particularly if stimulation-evoked latencies are comparable to previously-estimated delays in the response to sound in visual vs. auditory cortex). If, on the other hand, the initial visual response to sound largely results from subcortical transmission, the distribution of low-latency responses over occipital cortex should be comparable to what would be expected by chance alone (or to the profile produced by stimulation of a control region like the subcentral gyrus that is not expected to project to early visual cortex; Simonyan & Jürgens, 2002, 2003). In addition, to be consistent with nonhuman primate data, occipital N1 responses should be most prevalent/lowest-latency following stimulation of the mid/posterior portion of the superior temporal gyrus (STG), particularly at recording sites near V5/hMT+ and V1/V2 (and particularly over anterior portions of V1).

## Method

### Participants

Data were drawn from the open repository associated with van Blooijs et al., 2023 (data available at openneuro.org/datasets/ds004080). A subset (N = 23; mean age = 25 years, range 8-51; 11 female/12 male) of the original lifespan sample of 74 patients had combinations of STG stimulation and occipital recording sites that were relevant for the current question (see exclusion/inclusion criteria below), but which were not considered in the initial paper. In all cases, sites were positioned ipsilaterally, with the majority on the left hemisphere (17/23). For the positions of included stimulation pairs and recording sites for each patient, see Figure S1 (additional participant information can be found in Supplemental Table 1 and in the original repository).

#### Participant-level exclusion/inclusion criteria

Data were first screened to identify patients with combinations of STG and occipital coverage. Patients with relevant coverage were included if they had (a) at least one stimulation pair in which both electrodes were located on the STG, (b) at least one occipital recording location, and (c) at least 10 trials for analysis for at least one relevant stimulation pair.

### Data Acquisition

For details of data acquisition, readers are referred to van Blooijs et al. (2023). In brief, single-pulse electrical stimulation was conducted as iEEG data were recorded from subdural electrode strips/grids with 1 cm spacing. Data was initially acquired at a sampling rate of either 512 (n = 1) or 2048 Hz (n = 22). For a given pair of adjacent electrodes, stimulation (pulse width: 1ms, current intensity: 8mA) was typically applied up to 10 times at intervals of 5s, although in some cases larger numbers of stimulation pulses were recorded.

### Processing & Analysis

#### Selection of stimulation pairs and recording sites

Labels from the Destrieux et al. (2010) parcellation were used for initial identification of occipital, STG, and subcentral electrodes. The subcentral gyrus was selected as a control stimulation region given its comparable distance to occipital cortex and the expectation of limited communication between subcentral and visual areas (e.g. Simonyan & Jurgens, 2002; 2003). For delineation of regions within STG and occipital cortex, we used labels from the multimodal parcellation provided by Glasser et al. (2016; ‘HCP-MMP’) mapped to Freesurfer average space.

In SPES, electrical current is passed between a pair of electrodes, stimulating the cortex between them. Stimulation pairs were included in the following analyses if both members of a pair were located over the STG, or if one member was located on the STG and the other was located on the inferior portion of the supramarginal gyrus. The region ‘PSL’ from the HCP-MMP parcellation (Glasser et al., 2016) was used to define the upper border for inclusion of supramarginal electrodes. Pairs that crossed the superior temporal sulcus or lateral fissure were not included (to conservatively isolate stimulation to the STG, as far as possible). Control stimulation pairs were included if they were centered within the subcentral gyrus, excluding pairs that crossed the lateral fissure.

Stimulation pair/recording site combinations were also excluded if they contained any channels marked for exclusion by the criteria described in van Blooijs et al. (2023), or if the total number of trials was less than 10. Note: in a small set of cases (three patients, and 28 CCEPs total) trial counts were between 10 and 20; In all other cases, 10 trials contributed to each CCEP waveform. All trials for a given stimulation pair were included, except in rare instances in which stimulation occurred within 3s of another trial. Additional exclusion criteria based on data quality are described below.

#### Preprocessing and measurement of evoked responses

As in van Blooijs et al (2023), no modification of the continuous signal was performed (i.e., data were neither rereferenced from the original mastoid recording reference nor filtered). For measurement of evoked responses, timepoints between -12 and 12ms were excluded from analysis to limit contamination from stimulation artifacts (for a visualization of the range of the stimulation artifact see Figure S2). CCEPs were evaluated relative to a prestimulus average baseline of [-200 ms to -12 ms]. For peak latency measurement, the averaged response for a given combination of electrode site and stimulation pair was transformed by z-scoring the waveform relative to the mean and standard deviation over the baseline window. Peaks were identified as the first minimum/maximum in the interval [12-250ms] with an absolute value exceeding z = 5 and a minimum peak prominence of 2 (i.e. peaks were required to have a drop of z = 2 on either side before encountering a neighboring peak of greater amplitude). Electrodes were defined as showing a rapid response (N1) if the CCEP waveform contained a detectable peak in the range 12-80ms following stimulation (see Figures 2B and S3 for visualization of the set of CCEPs, sorted by latency). In the remainder of the document, the term CCEP is used to refer to the averaged waveform for a single stimulation pair/recording site combination.

**Figure 2.**
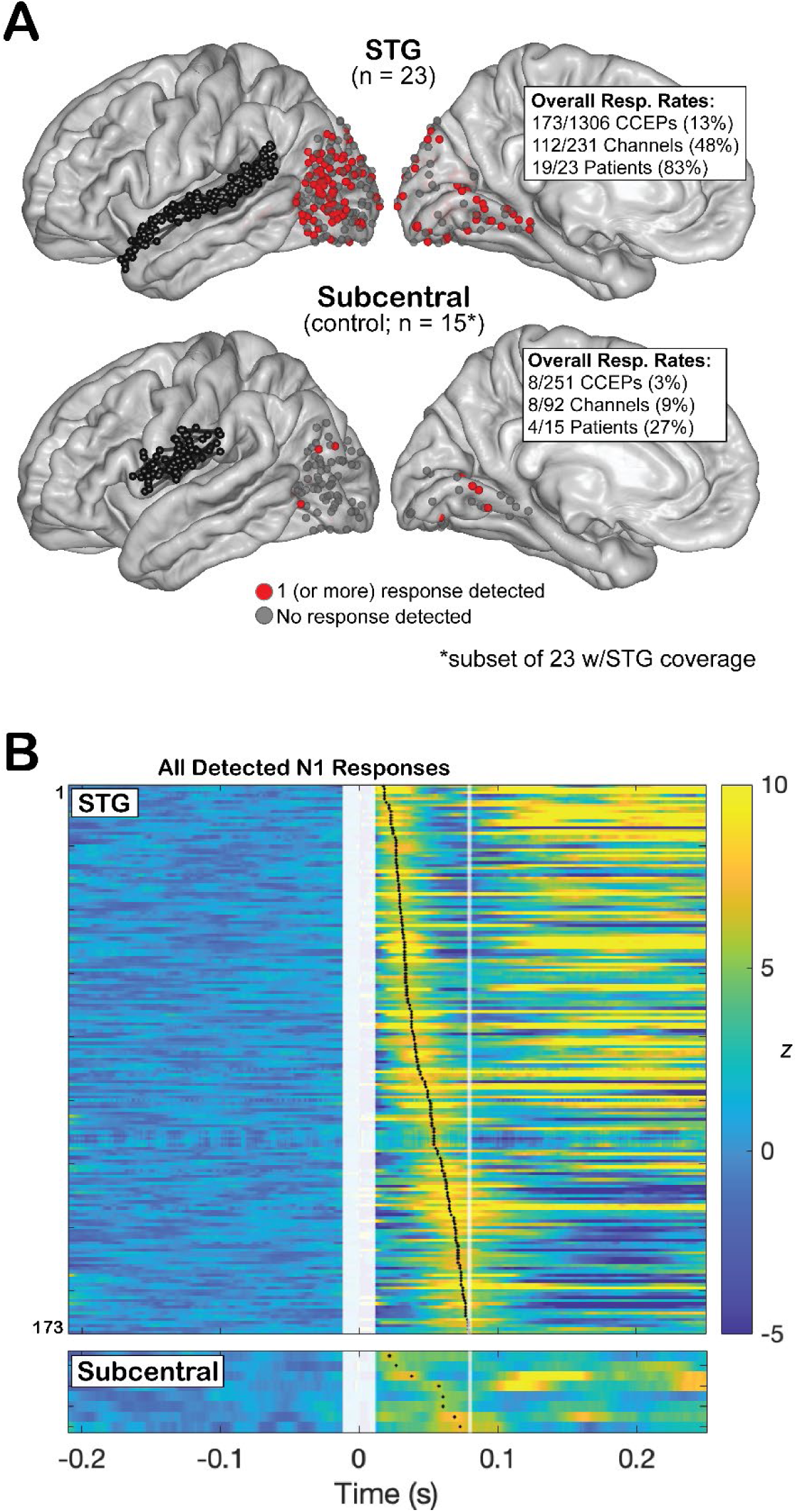
Summary of occipital responses to STG and Subcentral stimulation. (A) All stimulation pairs and recording sites for STG (upper) and Subcentral (lower) stimulation. Occipital recording sites are colored according to whether a response was observed at the site (red = at east one response detected; grey = no response detected). Note that the 15 patients with Subcentral coverage are a subset of the 23 with STG coverage. (B) Heatmap of all CCEP waveforms for which N1 responses were detected, sorted by atency (upper = STG stimulation; ower = subcentral stimulation). Z-scored responses were rectified such that the N1 peak was positive for all waveforms. The white bar surrounding timepoint 0 indicates the stimulation artifact. The white ine indicates the cutoff for N1 definition (80ms).

Saturated responses (defined by post-stimulation amplitudes exceeding 2000 microvolts) were excluded from analysis (n = 3 responses, 1 patient), as were CCEPs that showed extreme time-locked line-noise contamination (defined as having a mean power spectral density over the range 50-60 Hz, prior to stimulation, that was greater than 10x the median value over all stimulation pairs and electrode sites for that patient; n = 66 responses over 9 patients; this exclusion criterion was data-driven and implemented to reduce the occurrence of false-positive ‘peaks’ resulting from line noise). Overall, after these data quality exclusions, 1557 combinations of stimulation pair and recording site (1306 STG stimulation; 251 subcentral stimulation) were considered for analysis (231 total recording sites).

Given that the z-scored waveforms used to compute peak latency are scored relative to variation in the *averaged* waveform over the baseline window, the z-scores alone contain no information regarding variance in the waveform from trial to trial. To provide a coarse index of stability/variation in amplitude across repeated stimulations, peak latency values were supplemented with Cohen’s *d^z^* values computed for peak amplitude (computed by dividing peak amplitude by the standard deviation of the waveform across trials at that timepoint).

#### Spectral analyses

Spectral decomposition was performed to characterize the frequency content of the signal over all CCEPs in each region of interest. The intent of this decomposition was descriptive. Trials were epoched from -3 to 3s to allow a sufficient window for event-related spectral perturbation (ERSP) computation. The stimulation artifact was removed by interpolating the time-series between -12 and 12 ms for each trial with piecewise cubic hermite interpolation. Single-trial spectrograms were computed by convolving the iEEG time series with complex Morlet wavelets (for lower frequencies: 49 center frequencies linearly spaced between 2-50 Hz, with number of cycles per frequency increasing linearly from 3 to 12; for high-gamma power: 16 center frequencies linearly spaced between 75 and 150 Hz, with number of cycles per frequency increasing linearly from 12 to 24). A time step of ∼4ms was applied to produce time-frequency plots sampled at 256 Hz. Following convolution, power was extracted as squared amplitude for each trial, timepoint, and frequency. Spectrograms were baseline-corrected and normalized using the ‘vssum’ approach: for each timepoint the average power over time and trials in the interval [-500 to -200ms] at that frequency was subtracted, and values were then normalized by dividing by the sum of the value at that timepoint and the associated baseline value. The resulting values are bounded between -1 and 1 and scaled such that +/-0.33 indicates a 2x increase/decrease in power from baseline. High-gamma power values were averaged across all frequencies to produce a single time-series representation of high frequency activity.

### Data analysis

A nonparametric randomization approach was adopted to provide chance baselines against which to compare the observed data. To generate a baseline that best reflects the expected false-positive rate in the absence of stimulation, we generated a comparison distribution out of 1000 repetitions of the following: For each CCEP time-series, we extracted a matched null time-series centered around a point within the 2 seconds prior to a trial (selected randomly for each repetition; excluding baseline windows showing saturation or line noise contamination defined by the criteria outlined above). These null time-series were then processed in the same way as the observed data described above, resulting in 1000 surrogate datasets identical to the main dataset except for the use of null data. In this way, it was possible to evaluate expected false positive rates assuming no effect of stimulation for every observed statistic. Observed values were evaluated against this null distribution, and surrogate p-values were computed as the proportion of null values matching or exceeding the magnitude of the observed statistic. When observed values exceeded 100% of the null distribution, the surrogate p-value was recorded as “<.001” (the minimum observable p-value with 1000 datapoints). The 99th percentile of the null distribution was used as a benchmark for evaluating response rates in a given region.

#### Summary statistics by stimulation location and region

Stimulation pairs over STG were grouped by the HCP-MMP parcellation label associated with the nearest coordinate in Freesurfer average space to the center of the pair. Stimulation locations with limited representation were combined with neighboring locations to create four summary regions: posterior STG, mid-STG 1 (PBelt/A4), mid-STG 2 (A5), and anterior STG (Figure 1B). For region-of-interest (ROI) analyses over visual locations, we examined V1/V2, V3/V4, and V5/hMT+ (defined by HCP-MMP labels). Two of these regions, V1/V2 and hMT+, reflect hypothesized loci of early cross-modal input (Falchier et al., 2002; Majka et al., 2019; Palmer & Rosa, 2006; Rockland & Ojima, 2003) and were the subject of detailed examination; V3/V4 was included to provide a point of comparison. Sites were defined as near V5/hMT+ when the nearest coordinate in Freesurfer average space was positioned within the bounds of HCP-MMP regions MT/MST, TPOJ3/LO3, or V4t/FST (Figure 1B). These regions approximately cover the expected location of human motion-sensitive cortex (which varies from subject to subject) as defined by multiple atlases (see Huang et al., 2019 for a side-by-side visualization).

Patients with right-and left-hemispheric implants were first examined separately to evaluate whether there was any evidence of hemisphere-specific effects. No effects of hemisphere were identified, and data from both hemispheres were therefore pooled prior to analysis (Note, however, that ‘no effects of hemisphere’ should be understood in the context of sample size; we lack power in this sample to perform a detailed analysis of laterality effects).

To avoid pseudoreplication as far as possible, all response rates and latency values (barring initial summary statistics) were reported either in terms of the number of patients showing an effect, the effect per patient (i.e., one value per patient), or in the context of linear mixed effects (lme) models (using maximum likelihood estimation) accounting for patient identity (generated using the lme4 package in R; Bates et al., 2015). Models and summary statistics are described in the results section in each case. To provide estimates of uncertainty for fixed effects derived from lme models, confidence intervals (CI) around fixed effects (95%) were computed using the Wald method.

## Results

All data and code to reproduce the following analyses will be made available upon acceptance for publication. We examined occipital CCEPs following stimulation of STG in 23 patients, with a focus on the presence and latency of N1 responses occurring within 80ms of stimulation. Unless otherwise specified, *p*-values indicate the proportion of values computed from surrogate data that exceed the observed value. Overall, 173/1306 CCEPs exhibited detectable N1 responses to STG stimulation. At least one response was detected in 112/231 occipital sites, and at least one responsive site was observed in 19/23 patients (*p* = .003; Figures 2 & S3). The average response rate per patient was 17% (*sd* = 22%, *p* < .001). Responses to subcentral stimulation were comparatively limited, with 8/251 responses detected over 8/92 occipital sites across 4/15 patients (*p* = .914). The average response rate to subcentral stimulation per patient was 3% (sd = 8%; *p* = .495).

Figure 3 summarizes occipital N1 responses as a function of stimulation location along the STG. The average response rate across patients was greatest following stimulation of posterior STG, and declined with increasingly anterior stimulation (Figure 3B; Average response rates and surrogate p-values: Anterior STG: *M_resp_rate_* = .10, *p* = .003; Mid STG 2: *M_resp_rate_* = .11, *p* = .001; Mid STG 1: *M_resp_rate_* = .14, *p* < .001; Posterior STG: *M_resp_rate_* = .25, *p* < .001; compare with subcentral response rates described above). Qualitatively, N1 responses to posterior STG stimulation tended to concentrate around lateral occipital cortex, becoming more diffuse (and longer-latency) as stimulation approached anterior STG (Figure 3A).

**Figure 3.**
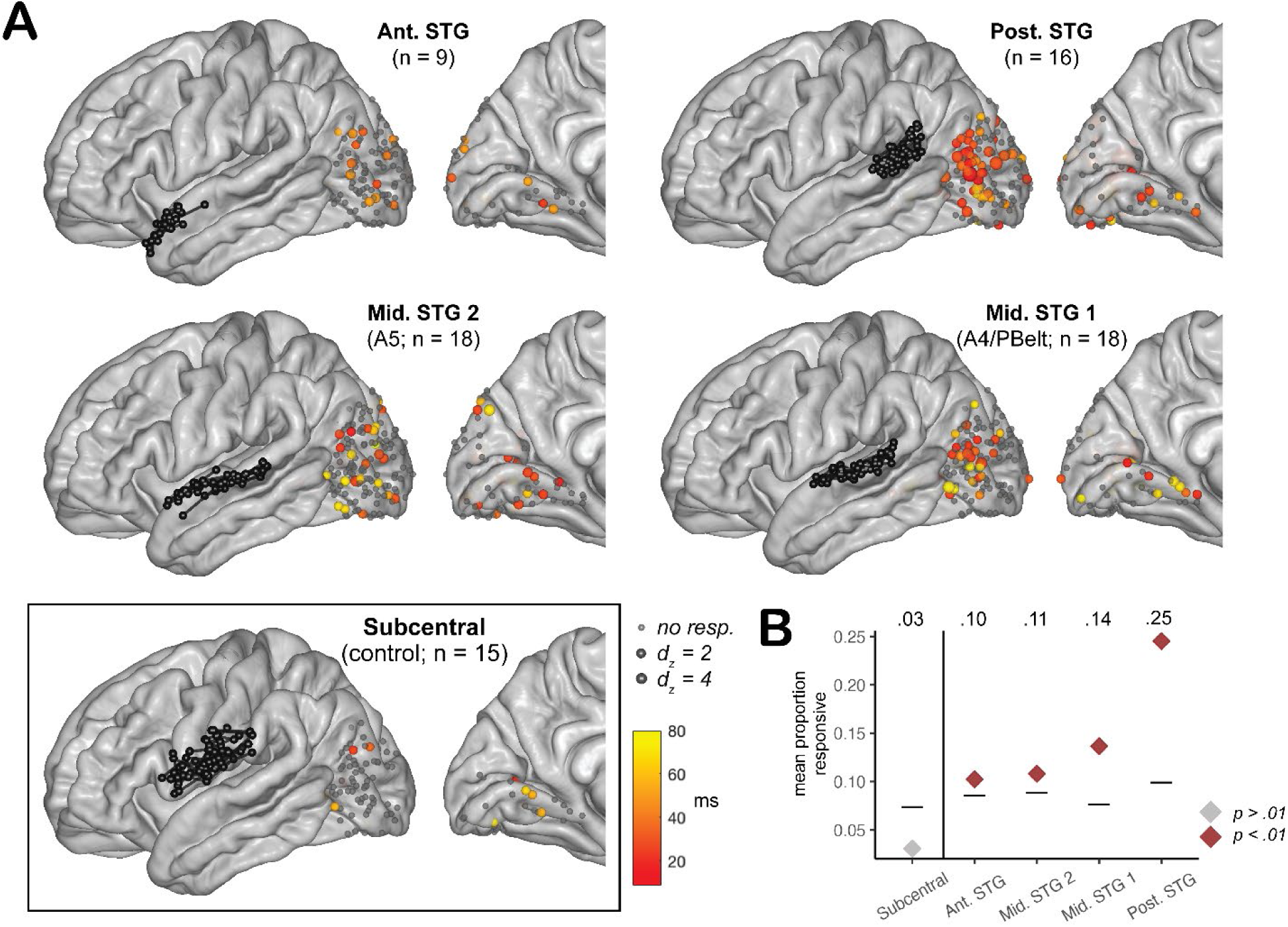
Distribution and latency of occipital responses to STG and Subcentral stimulation. (A) Stimulation pairs and recording sites presented separately for each stimulation ROI. Each recording site is colored according to the fastest detected response at that location (between 12-80ms). Grey indicates no response was detected at that site. The size of each recording site is scaled based on the standardized effect size (Cohen’s d_z_) for the corresponding response. (B) Average response rate across patients for each stimulation ROI. Color indicates whether the response rate exceeded chance baseline (99^th^ percentile) derived from null data. Black lines indicate the 99^th^ percentile of the null distribution.

To characterize the distribution of responses, we report response rates and N1 latencies as a function of stimulation location and occipital ROI in Figure 4. Response rates were first evaluated against chance models *within* each stimulation/recording region combination, and then compared across all pairs of STG stimulation regions with mixed-effects models that included a random effect of patient identity and a fixed effect of stimulation location (surrogate p-values for effects of stimulation location were derived by fitting identical models to the null data and comparing the observed fixed effect against the distribution of null fixed effects; FDR correction was performed across all comparisons using the Benjamini-Hochberg approach). Within V5/hMT+, both mid- and posterior-STG stimulation yielded above-chance response rates (Figure 4A). The proportion of detected responses per patient increased with proximity as stimulation moved from anterior (5%) to posterior STG (28%; Figure 4A; FDR-corrected p < .05 for all contrasts that included Posterior STG; all other contrasts *n.s.*). Within V1/V2, response rates only exceeded chance levels following mid-STG stimulation (i.e., when stimulation was centered over likely auditory association areas accessible to contacts placed on the outer surface of mid-STG), and declined with more anterior/posterior stimulation (Figure 4A; note that only the reduction in response rate between region “Mid. STG 1” and Posterior STG was robust to FDR correction; all other contrasts *n.s.*). This effect appears largely driven by responses in anterior portions of the calcarine sulcus and lingual gyrus (Figure 3A). Within V3/V4, response rates did not show clear differentiation by stimulation location.

**Figure 4.**
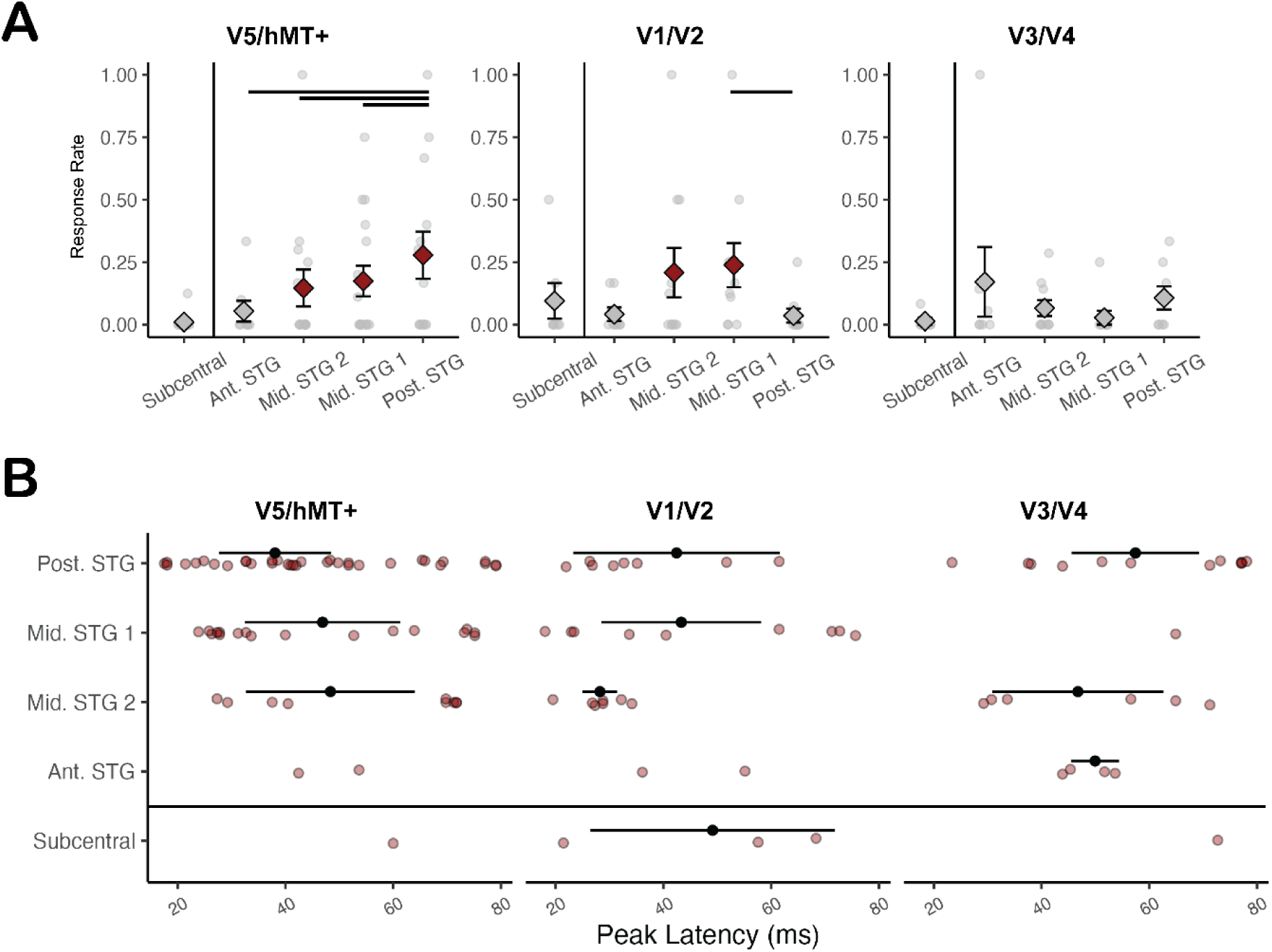
Response rates and latencies for each occipital ROI. (A) Response rates by stimulation location. Each grey dot indicates the proportion of responsive CCEPs relative to the total number of CCEP waveforms for a single patient. Diamonds indicate the average response rate across patients (error bars = SEM). Red diamonds indicate situations in which the average response rate exceeded the 99^th^ percentile of the associated chance baseline (as in Figure 3). Horizontal bars indicate groups for which differences in response rates were greater than expected given the chance distribution derived from null data (after FDR correction). (B) Latencies for all detected N1 responses (red points). Estimates of mean latency and associated 95% CIs derived from LME models are plotted with black points/bars (where sufficient data was available).

Latencies of all detected N1 responses are presented by ROI in Figure 4B and summarized in Table 1 (along with 95% CIs and latencies of the fastest observed response in each case). Latencies for each combination of stimulation/recording ROI were summarized (when sufficient data was available) with simple intercept-only lme models including a random effect of patient identity. The fixed intercept of each model and associated 95% CI were used to estimate population mean latency (Bates et al., 2015). Given that there were pronounced differences in response rate as a function of stimulation location, the following description of latency distributions should be interpreted with caution. Within V5/hMT+, the distribution of responses to mid-STG stimulation (both regions) was somewhat bimodal, with most N1 peaks hovering between ∼25-40ms, and a few outlying responses between 70-80ms, resulting in overall latency estimates of *β* = 47-48ms; Responses to posterior STG stimulation were also broadly distributed, but with a somewhat greater concentration of low-latency responses (<40ms), resulting in an estimated overall latency of *β* = 38ms (10ms faster on average than mid-STG stimulation; Figure 4B; Table 1). The fastest overall response in V5/hMT+ (18ms) was also recorded following stimulation of posterior STG. Anterior STG and subcentral stimulation yielded too few suprathreshold responses to derive reasonable estimates of latency, and all detected responses were >40ms.

**Table 1.**
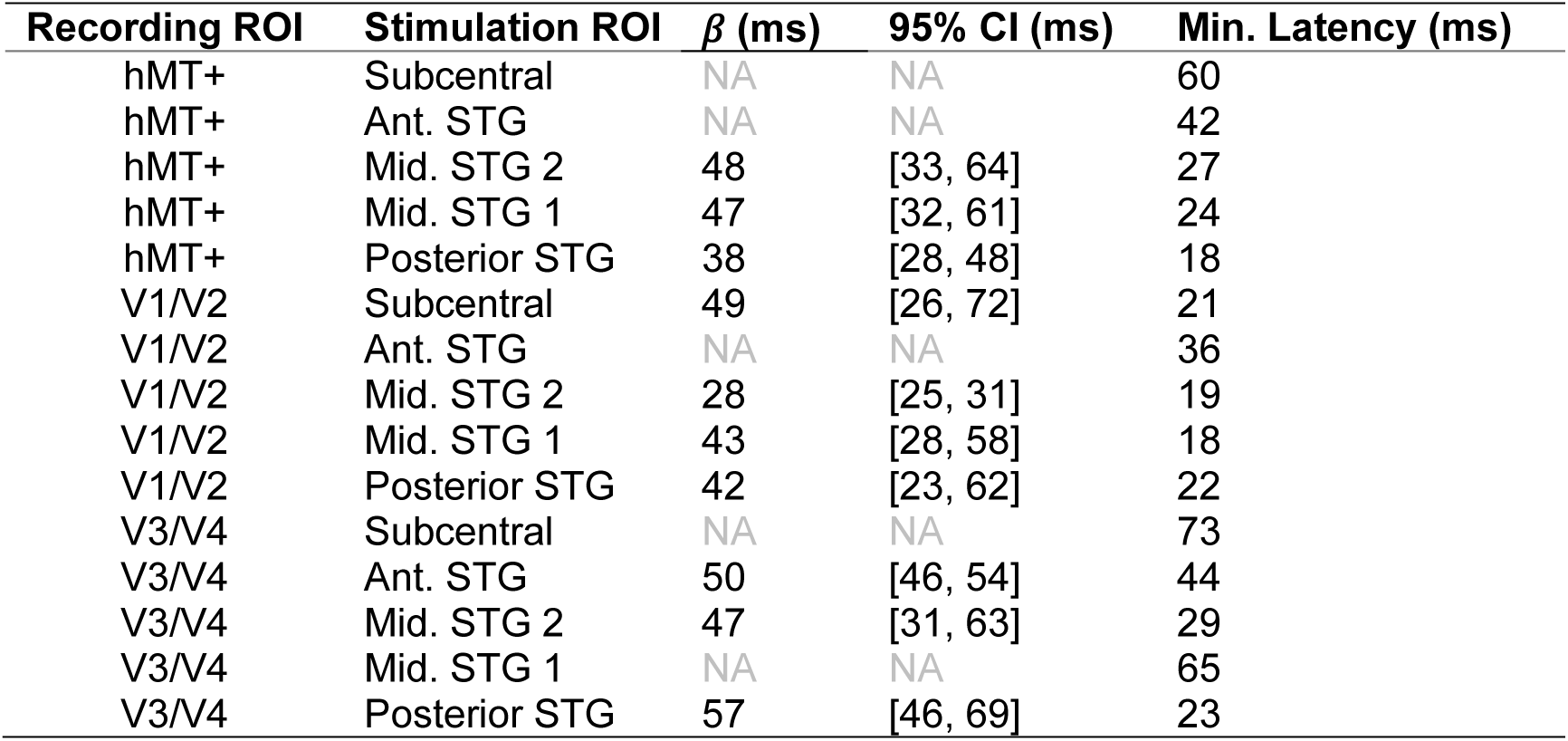
N1 Latency estimates by stimulation and recording ROI.

| Recording ROI | Stimulation ROI | $\beta$ (ms) | 95% CI (ms) | Min. Latency (ms) |
| --- | --- | --- | --- | --- |
| hMT+ | Subcentral | NA | NA | 60 |
| hMT+ | Ant. STG | NA | NA | 42 |
| hMT+ | Mid. STG 2 | 48 | [33, 64] | 27 |
| hMT+ | Mid. STG 1 | 47 | [32, 61] | 24 |
| hMT+ | Posterior STG | 38 | [28, 48] | 18 |
| V1/V2 | Subcentral | 49 | [26, 72] | 21 |
| V1/V2 | Ant. STG | NA | NA | 36 |
| V1/V2 | Mid. STG 2 | 28 | [25, 31] | 19 |
| V1/V2 | Mid. STG 1 | 43 | [28, 58] | 18 |
| V1/V2 | Posterior STG | 42 | [23, 62] | 22 |
| V3/V4 | Subcentral | NA | NA | 73 |
| V3/V4 | Ant. STG | 50 | [46, 54] | 44 |
| V3/V4 | Mid. STG 2 | 47 | [31, 63] | 29 |
| V3/V4 | Mid. STG 1 | NA | NA | 65 |
| V3/V4 | Posterior STG | 57 | [46, 69] | 23 |

Within V1/V2, there were sufficient numbers of detected N1 responses to derive latency estimates for posterior STG and both mid-STG locations. When stimulation occurred over posterior STG or its neighboring mid-STG ROI (Mid. STG 1; corresponding to areas A4/PBelt of the HCP-MMP atlas), the distribution of latencies was fairly broad, producing latency estimates of *β =* 42-43ms (although a number of peaks in both cases emerged before 40ms). Responses to stimulation of the more anterior mid-STG ROI (Mid. STG 2; corresponding to area A5 of the HCP-MMP atlas) all emerged before 40ms (*β = 28ms*), deviating somewhat from the more distributed latencies observed for other combinations of stimulation/recording site. The fastest latencies in V1/V2 (18-19ms) emerged following stimulation of both mid-STG regions. As with V5/hMT+, anterior STG stimulation produced too few detectable responses to generate a latency estimate (and the responses that were detected were comparatively slow; Figure 4B). Subcentral stimulation also yielded few responses, with latencies that were widely distributed.

Within V3/V4, response latencies were heterogeneous. Although some low-latency responses were observed, across all stimulation regions, latency estimates hovered between *β =* 47-57ms.

The spectral content of CCEPs was qualitatively similar from region to region (Figures S5 and S6). In both hMT+ and V1/V2, CCEPs were largely dominated by low-frequency activity, with no evidence of stimulation-evoked activity in the high-gamma range. These patterns are consistent with modulatory effects produced by feedback-type signal propagation, with no indication of stimulation-elicited local neuronal spiking typically indexed by increased high-frequency activity (Bastos et al., 2015; Buffalo et al., 2011). Such patterns map well to the anatomy of primate corticocortical projections, which also have a feedback/modulatory character (Falchier et al., 2012).

### Exploratory analyses of V1 ROI

In nonhuman primates, auditory projections preferentially target anterior regions of V1 corresponding to representation of the peripheral visual field (Falchier et al., 2002; Rockland & Ojima, 2003). In this dataset, electrode coverage of multiple locations along V1 provided a rare opportunity to explore the presence of a similar bias in the human CCEP. To this end, we examined response rates and latencies as a function of the position of V1 recording sites along the anterior-posterior axis (coarsely grouped into anterior/mid/posterior locations by Freesurfer average y-coordinate). As shown in Figures 5 & 6, responses were detected in the most anterior portion of the calcarine sulcus (with an average detection rate of 33% across 3 patients with anterior calcarine sites), and to some extent the occipital pole (9% across 4 patients; below the corresponding chance threshold). There were no detectable N1 responses in mid-calcarine locations typically associated with parafoveal processing (Figure 5; Note: to supplement the grouped data, we also performed a sliding-window assessment of response rate, grouping CCEPs by y-coordinate with a step of 5 and a window size of 10; Results using this approach corroborate the grouped data; Figure 5B). The latency of anterior visual responses ranged from 18-61ms (Figure 5D), with 5/8 peaks emerging within 30ms of stimulation (β = 33 ms, 95% CI [22, 45]). The latency of occipital polar responses ranged from 34-76ms (β = 45 ms, 95% CI [29, 62]).

**Figure 5.**
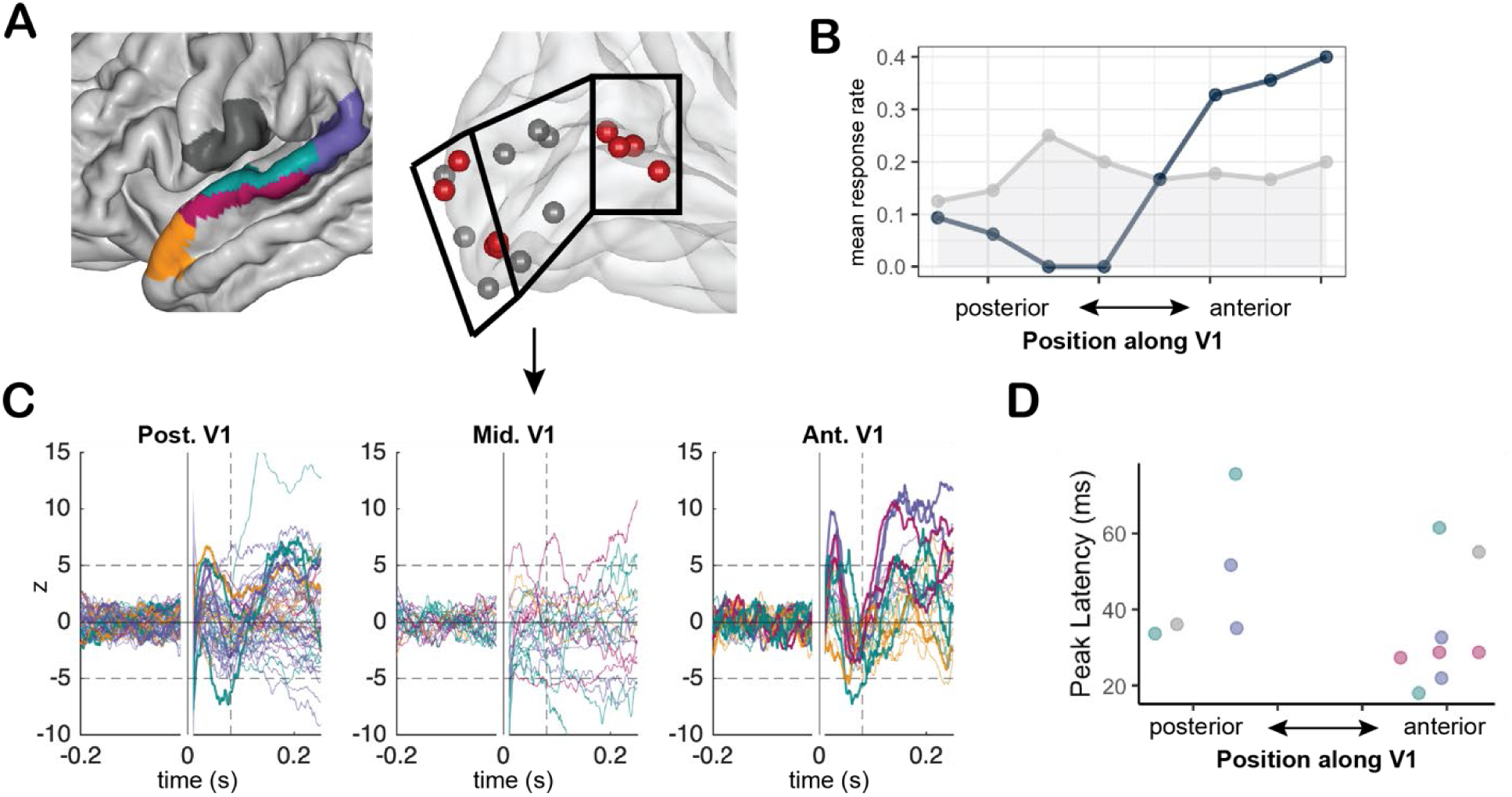
V1 responses by eccentricity. (A) Summary of responsive (red) and nonresponsive (grey) V1 sites. (B) Average response rate as a function of position along V1 (from posterior to anterior). Dark line indicates observed response rate; Grey shaded region indicates 99^th^ percentile of the permutation distribution at each eccentricity. (C) All CCEP waveforms, colored according to stimulation region as shown in A, and split by location along V1. Dashed vertical/horizontal lines indicate the thresholds for response detection. (D) Latencies of all observed N1 responses, plotted by position along V1. Color indicates stimulation location by the key shown in A.

**Figure 6.**
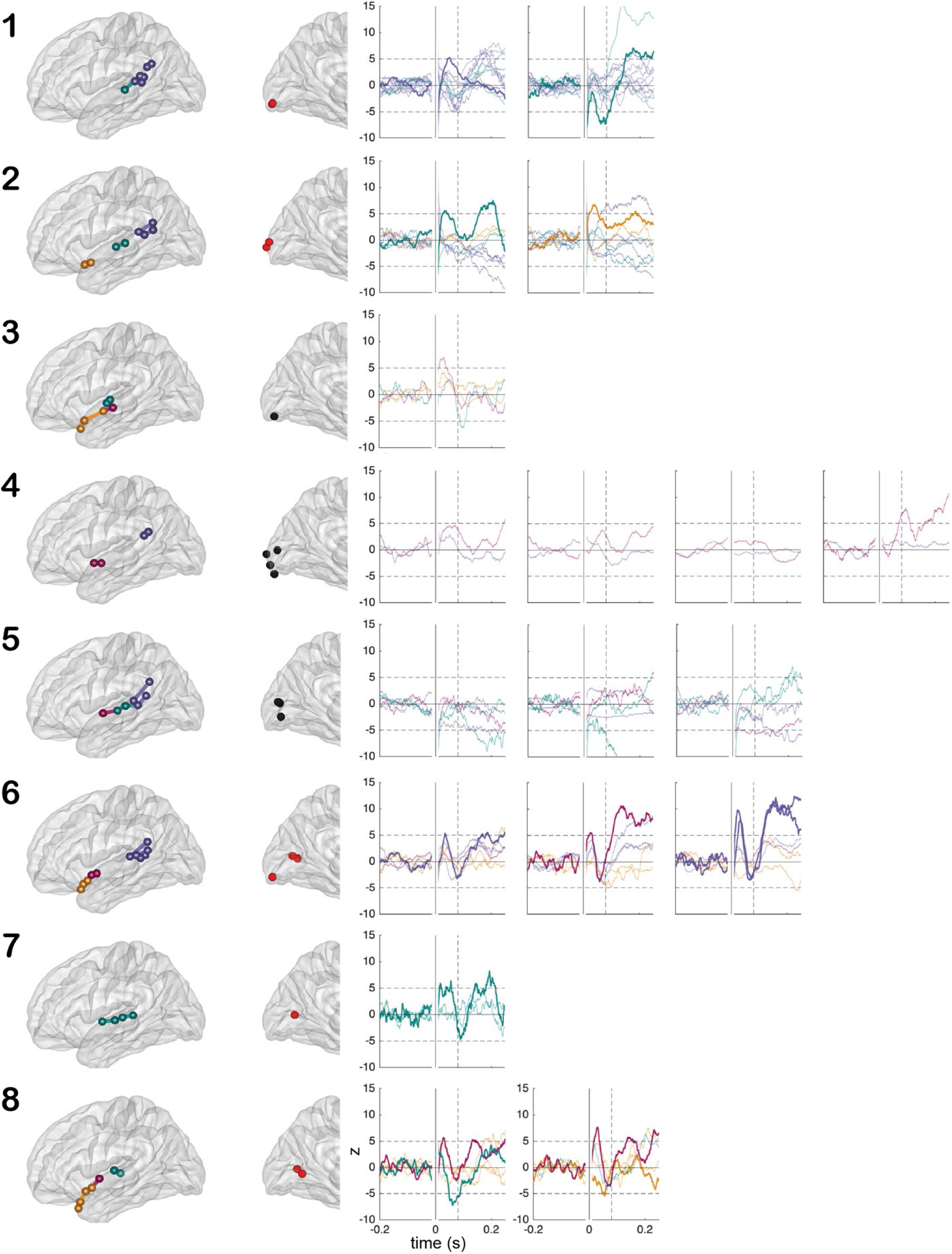
CCEP waveforms for all patients with V1 recording sites. Each row contains data for a single patient, ordered from top to bottom according to approximate location of recording sites along V1. Stimulation pairs are colored according to stimulation region. Recording sites are colored according to whether a response was detected at that site. Each waveform plot contains data from a single recording site (with waveforms colored according to stimulation region). Waveform plots for each patient are arranged left-to-right by location of the recording site (anterior-posterior). Waveforms with detected N1 responses are plotted in “bold.”

### Exploratory analyses of V5/hMT+ ROI

Reports of sound-evoked visual activity (Plass et al., 2019) and human/nonhuman primate connectivity (Gurtubay-Antolin et al., 2021; Majka et al., 2019; Palmer & Rosa, 2006) also highlight V5/hMT+ as a likely site of early cross-modal input, particularly from auditory motion-sensitive areas along the planum temporale. This is consistent with the observation of posterior-STG-dominant responses in the broad V5/hMT+ ROI (Figure 1D). To clarify the specificity of responses within this region, we examined CCEPs within three subregions defining the greater ROI (MT/MST, the more dorsal area covered by LO3/TPOJ3, and the more ventral area V4t/FST). Although there were a larger number of detected responses overall in sites located within LO3/TPOJ3, the response to posterior STG stimulation was maintained across all three locations (Figure 7). Notably, within MT/MST, responses to stimulation from other locations along the STG were comparatively limited (that is, responses in MT/MST showed some selectivity to posterior STG; Figure 7B). With respect to latency, estimates in MT/MST and LO3/TPOJ3 following stimulation of posterior STG were roughly comparable (*β*s = 37-38ms; though the distribution of responses in LO3/TPOJ3 was notably denser, with a larger number of low-latency responses to both mid- and posterior STG stimulation; Figure 7C). Responses in V4t/FST tended to emerge later (*_β_*_post_STG_ = 57ms), consistent with a dorsal bias in communication from auditory areas (see Supplemental Table 2 for a full summary of latency estimates by stimulation location for each hMT+ subregion).

**Figure 7.**
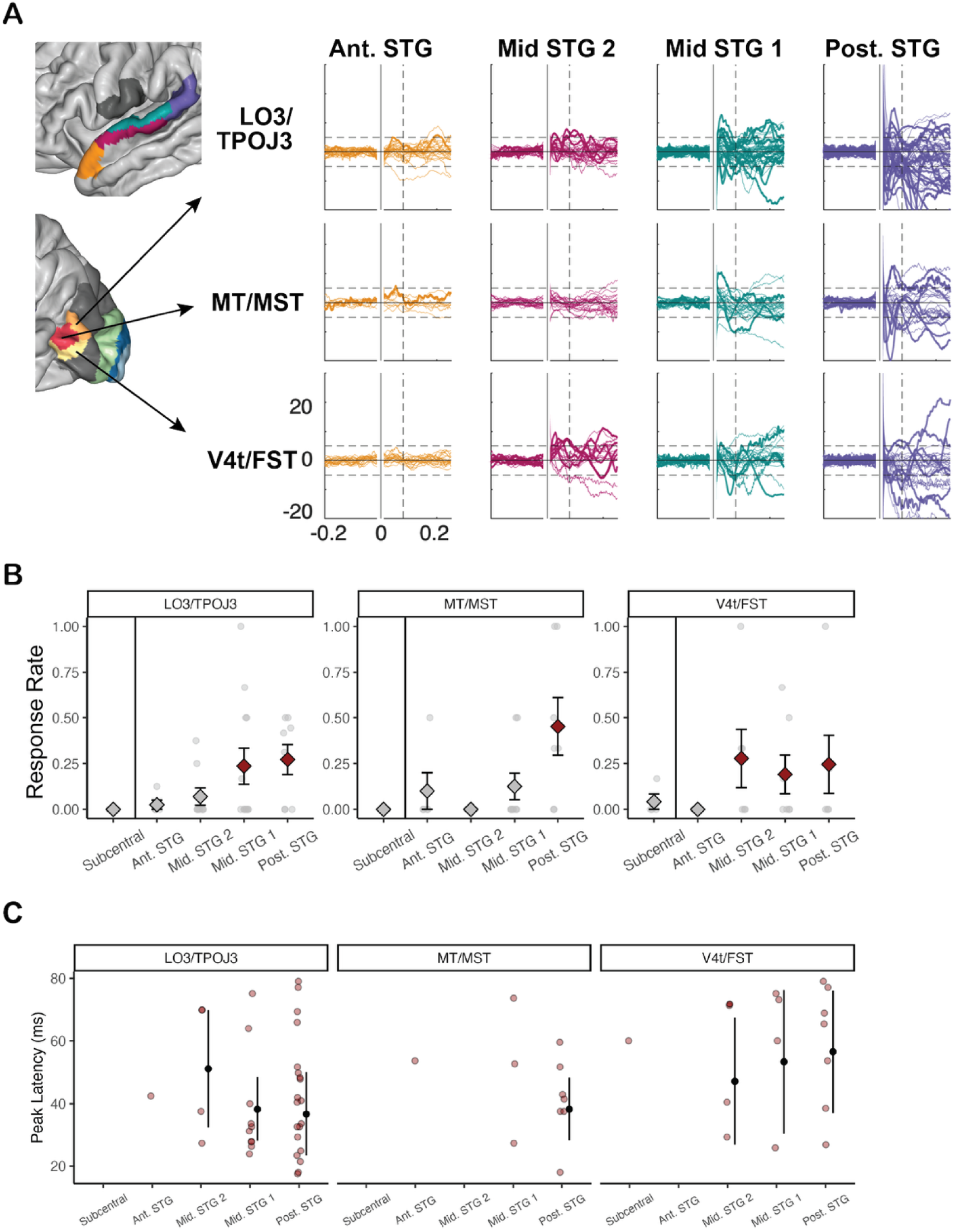
Summary of hMT+ responses by subregion. (A) All CCEPs for all three subregions of the greater V5/hMT+ ROI, separated by stimulation location along STG. Dashed vertical/horizontal lines indicate thresholds for N1 evaluation. (B) Average response rates for each subregion, plotted in the same manner as Figure 4. (C) Response latencies for all N1 responses in each subregion, plotted in the same manner as Figure 4.

## Discussion

An accumulating body of evidence indirectly supports the rapid transmission of information from auditory to visual cortex in humans. Here, we extend these observations by characterizing effective connectivity between superior temporal and occipital cortex in the context of intracranial recordings during single-pulse electrical stimulation in 23 patients with epilepsy (van Blooijs et al., 2023).

Early visual cortex responded rapidly (18ms+) to electrical stimulation over mid- and posterior STG, with a distribution that largely matched patterns of connectivity observed in nonhuman primates (Falchier et al., 2002; Rockland & Ojima, 2003) and non-invasive work in humans (Beer et al., 2011; Gurtubay-Antolin et al., 2021). That is, N1 responses were most prevalent in lateral occipital cortex, particularly following stimulation of the posterior portion of the STG, and showed a degree of bias toward the anterior portion of V1. Although functional localizers were not available for these patients, the distribution of stimulation-evoked responses also corresponded well to reports of sound-evoked responses in peripheral V1 (Brang et al., 2015; Plass et al., 2019; Ferraro et al., 2020), and reports of auditory motion-related activation of V5/hMT+ and adjacent cortex (Rezk et al., 2020). The current dataset contained only ipsilateral stimulation/recording combinations, but we expect that cross-modal corticocortical pathways between early sensory cortices are restricted to the same hemisphere, given animal work and prior research in humans suggesting primacy for spatially-aligned auditory-visual stimuli (McDonald et al., 2000; Plass et al., 2019; Wallace et al., 1996).

Animal studies suggest that auditory inputs to visual cortical neurons have a feedback-type laminar profile (Falchier et al., 2012) and exert a modulatory influence on visual cortical processing (Allman & Meredith, 2007; Meredith et al., 2009; Wang et al., 2008). That is, rather than directly triggering neural activity in visual cortex, auditory inputs may instead produce subthreshold changes in membrane potential, resulting in a phase-reset of ongoing oscillatory activity that facilitates the response to upcoming visual input (Mercier et al., 2013; Thorne & Debener, 2014). The spectral profiles of occipital CCEPs were broadly consistent with this type of effect. Superior temporal stimulation produced low-frequency modulation (<30 Hz; typical of feedback signal propagation; e.g. Bastos et al., 2015), without inducing the high-frequency response typically associated with neuronal spiking.

### V1 Responses

As summarized above, within V1, superior temporal stimulation evoked particularly strong N1 responses at anterior electrode sites. This distribution is consistent with nonhuman primate data, but to map well to the human sound-evoked visual response, the latencies of these CCEP responses should be comparable to observed delays between sound-evoked responses in auditory and visual cortex. The fastest N1 response in anterior V1 peaked at 18ms, and the average response latency was approximately 33ms. In comparison, visual responses to sound typically onset 2-15ms after the earliest detected response in auditory cortex (Brang et al., 2015; Raij et al., 2010). Although a response time of 18ms is broadly consistent with this sound-evoked data, one might argue that communication latencies near 30ms are too slow to account for the speed of the initial sound-evoked visual response. In our view, there are several ways to reconcile these observations. First, it is certainly possible that some portion of the initial sound-evoked response in occipital cortex results from subcortical rather than corticocortical transmission.However, it is worth noting that the approach to quantifying early responses used here is based on identification of peaks in the signal, whereas prior investigations of sound-evoked patterns have focused on response onset. In CCEP contexts, both very-low-latency peaks (<12ms) and onset values can be difficult to determine given that they are confounded to a degree with the stimulation artifact, but there is at least one large-N CCEP study in which N1 onset estimates reliably precede peak latency estimates by at least 10ms (Trebaul et al., 2018). Adjusting our peak latency estimates downward by ∼10ms would place many of the anterior calcarine observations in this study within approximately the same latency range as the delay in sound-evoked onset between auditory and visual cortex.

Notably, anterior calcarine responses were not constrained to a single stimulation region, but emerged following stimulation of both middle and posterior portions of the STG. The distribution of effects within the middle portion of the STG, in particular, is somewhat counterintuitive. Based on the nonhuman primate literature, in which auditory projections to peripheral V1 are most prevalent from the caudal belt and parabelt (Falchier et al., 2002; Rockland & Ojima, 2003), we expected to see strong anterior visual responses with stimulation between the transverse gyrus and planum temporale (e.g. when contacts were placed as in Figure 6, row 7). However, this was not the case, at least in our limited sample. V1 responses to stimulation of this region were relatively limited, and did not conform to the stereotypical CCEP N1/N2 shape. For example, in at least two cases (Figure 6, rows 2 and 7, green waveforms), the response was almost periodic, with wide, rounded peaks rather than sharp initial deflections. In addition, stimulation just anterior to the transverse gyrus yielded faster and more stereotypically-shaped initial peaks (Figure 6).

Interpretation of these observations relative to the literature is complicated not only by the sample size, but also by the placement of electrodes on the cortical surface in this dataset. This placement limits access to core and belt auditory regions embedded in Heschl’s gyrus, such that we can make only limited assertions regarding stimulation of early auditory cortex. To more completely describe communication between primary sensory cortices (and to better understand the results above), it will be necessary to assemble data in which stimulation is applied over contacts more directly adjacent to auditory belt/parabelt regions where many of the visual projections identified in macaques originate (Falchier et al., 2002, 2012; Rockland & Ojima, 2003). Such data would be particularly useful for comparison to sound-evoked visual responses, given that such responses may reflect communication from auditory cortical sources inaccessible to external cortical stimulation.

STG stimulation also evoked possible responses over the occipital pole, although there were no detectable responses in mid-calcarine sites. Occipital polar responses were generally weaker than those over the anterior calcarine sulcus (see Figure 6, subject 6), and they have limited support in nonhuman primate data, where the rate of auditory projections to V1 declines monotonically with decreasing visual eccentricity (approaching zero near the center of the visual field; Falchier et al., 2002). At least two intracranial reports in humans also describe limited, if any, response to sound in posterior occipital contacts (Mercier et al., 2013; Plass et al., 2019). However, such responses are compatible with results from human DTI that identify possible connections between auditory areas and both anterior and polar portions of early visual cortex (Beer et al., 2011), as well as cross-modal effects reported at the occipital pole (Romei et al., 2007, 2009), and could point to a bimodal distribution of projections in humans that diverges from that observed in nonhuman primates. Given that few patients had coverage of multiple locations within V1 in this sample, additional study will be necessary to thoroughly address this question.

The observation of responses near the occipital pole may also be consistent with reports that have used noninvasive scalp EEG to examine cross-sensory interactions. For example, combining an auditory stimulus with a visual stimulus will produce modulations of the parieto-occipital evoked response approximately 40-90ms after stimulation onset (Giard & Peronnet, 1999; Mishra et al., 2007; Molholm et al., 2002). This response has been linked to modulation of the C1 component (Molholm et al., 2002) thought to originate in striate cortex. It is difficult to imagine that a modulation of the C1 component observable at the scalp could arise from auditory inputs deep within the calcarine fissure. However, projections from auditory cortex to posterior calcarine areas (with a latency of ∼45ms) could plausibly yield such an interaction.

Notably, these noninvasive studies also report an asymmetry in modulation strength, with a bias toward the right hemisphere (Giard & Peronnet, 1999; Molholm et al., 2002). In our initial assessment of laterality effects (prior to pooling data across hemispheres), we did not observe differences as a function of hemisphere of stimulation. However, we also did not have sufficient statistical power in this sample to perform an extensive comparison of effects of stimulation across hemispheres, nor did we have any patients in this sample with concurrent recording in both hemispheres (which would allow within-patient investigation of asymmetric organization). The question of hemispheric asymmetry, like the question of contralateral vs. ipsilateral communication, remains an open avenue for inquiry.

### hMT+ Responses

The most pronounced responses to stimulation emerged over lateral occipital cortex, particularly following stimulation of the posterior STG. This pattern is not entirely surprising, given that lateral occipital cortex is closest in proximity to the sites of stimulation, and it is consistent with at least one prior report detailing the distribution of sound-evoked visual responses across human visual cortex (Plass et al., 2019). As proposed in Plass et al. (2019), these lateral clusters of responses may reflect communication from auditory cortex to visual motion-sensitive area V5/hMT+.

Although motion-based localizers were not available in this study, responses within anatomically-defined V5/hMT+ were compatible with this hypothesis. Within this region, responses were generally strongest following posterior STG stimulation, with a dorsal bias such that sites over MT/MST and neighboring LO3/TPOJ3 responded faster and more frequently than sites over V4t/FST. The earliest CCEP responses in MT/MST and LO3/TPOJ3 emerged ∼18ms following posterior STG stimulation, and the average latency of responses was approximately 38ms. To our knowledge, there are no existing estimates of relative latencies derived from concurrent recordings in planum temporale and hMT+ during auditory stimulation. However, in lateral occipital cortex, responses to sound tend to onset between 50-100ms following stimulation (Plass et al., 2019). In the planum temporale, initial auditory evoked potentials emerge around 30ms (Liégeois-Chauvel et al., 1994). Taken together, these reports suggest an approximate delay of 20ms between the earliest responses in both regions.

This pattern is consistent with the notion of a relay conveying auditory positional or motion-related information from posterior STG to visual motion-sensitive cortex. There is some support for the existence of corticocortical projections between auditory cortex/planum temporale and V5/hMT+ in both nonhuman primates (Majka et al., 2019; Palmer & Rosa, 2006) and humans (DTI: Gurtubay-Antolin et al., 2021). These observations are paralleled by functional studies demonstrating auditory motion-related responses in V5/hMT+ in the blind and, to a lesser extent, the sighted (Bedny et al., 2010; Dormal et al., 2016; Poirier et al., 2005, 2006; Rezk et al., 2020; Saenz et al., 2008; Strnad et al., 2013).

Studies of auditory motion in the blind/sighted highlight the potential value in comparing distributions of CCEP responses with patterns of sound-related visual recruitment following vision loss. That is, the profile of CCEP responses may provide insight into patterns of plasticity following disease/injury by illustrating where cross-modal corticocortical inputs in the sighted are most prevalent. This is likely most relevant for the late blind, where the distribution of projections is assumed to be similar at onset of vision loss to that of the sighted. However, our results also bear similarities to sound-evoked responses in the occipital cortex of early and congenitally blind individuals. For example, patterns of auditory scene decoding in the early visual cortex of congenitally blind participants also show a pronounced eccentricity bias (present to a lesser extent in sighted subjects; Vetter et al., 2020), and early blindness is associated with greater recruitment of visual motion-sensitive cortex in response to auditory motion (Poirier et al., 2006; Strnad et al., 2013; Bedny et al., 2010).

### Conclusions

Overall, these results establish that rapid communication between auditory and visual cortex can occur in humans and suggest that such communication is organized in a manner similar to that of nonhuman primates. The results are also consistent with the hypothesis that the initial response to sound in human visual cortex originates, at least in part, from auditory cortex. Importantly, these data cannot be used to rule out contributions of other pathways to the sound-evoked visual response (including subcortical, corticothalamocortical, and/or pathways through higher-order multisensory cortex). The respective contributions of these pathways remain an open avenue for inquiry. Although the current data was obtained from patients with epilepsy, the compatibility of our observations with reports of sound-evoked responses in other patients (Brang et al., 2015; Ferraro et al., 2020; Mercier et al., 2013; Plass et al., 2019), connectivity in nonpatient samples (Beer et al., 2011; Gurtubay-Antolin et al., 2021), and tractography in nonhuman primates (Falchier et al., 2012; Majka et al., 2019; Palmer & Rosa, 2006; Rockland & Ojima, 2003) provides a measure of confidence in the results.

## Supplemental Information

**Table S1.**
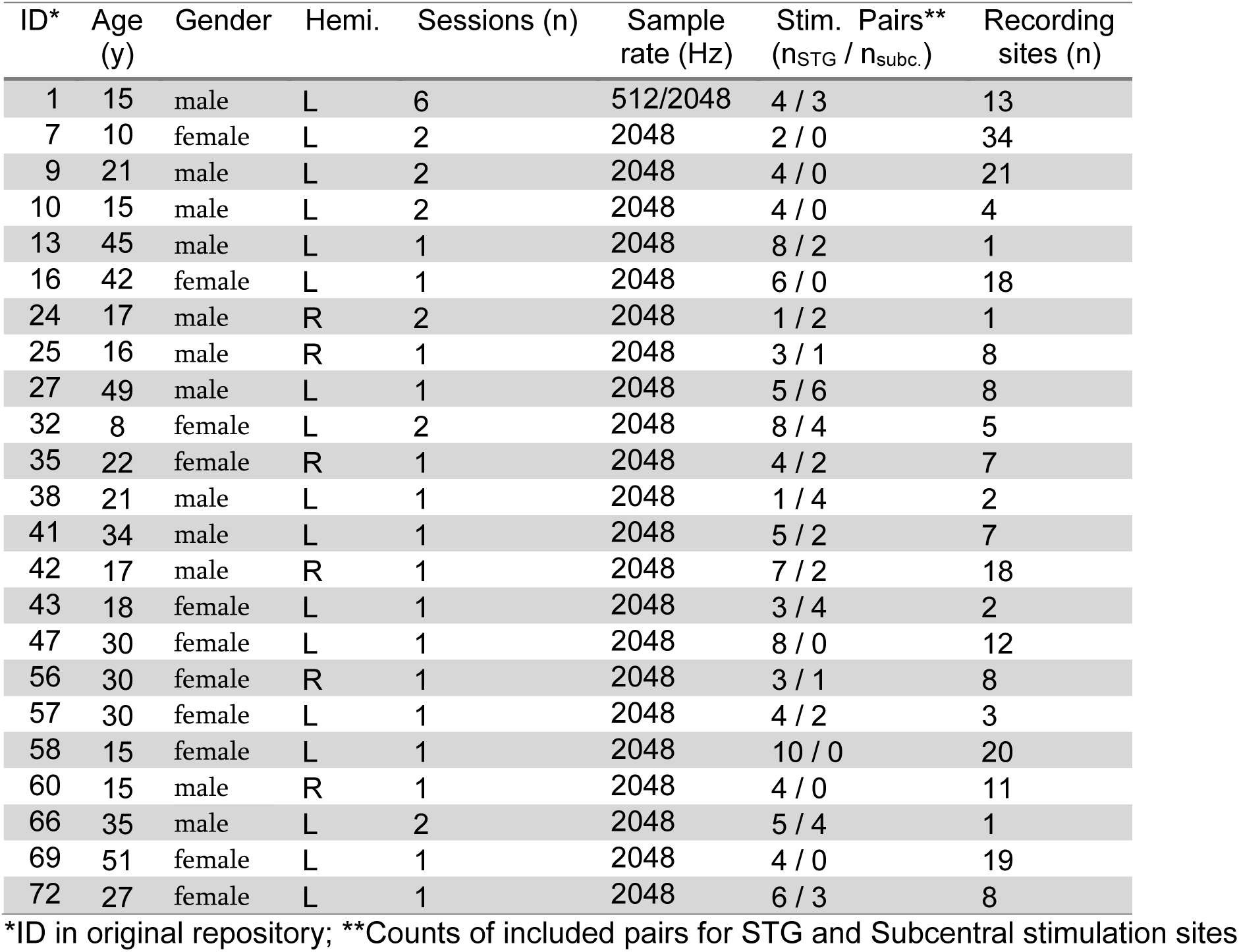
Patient details.

**Table S2.**
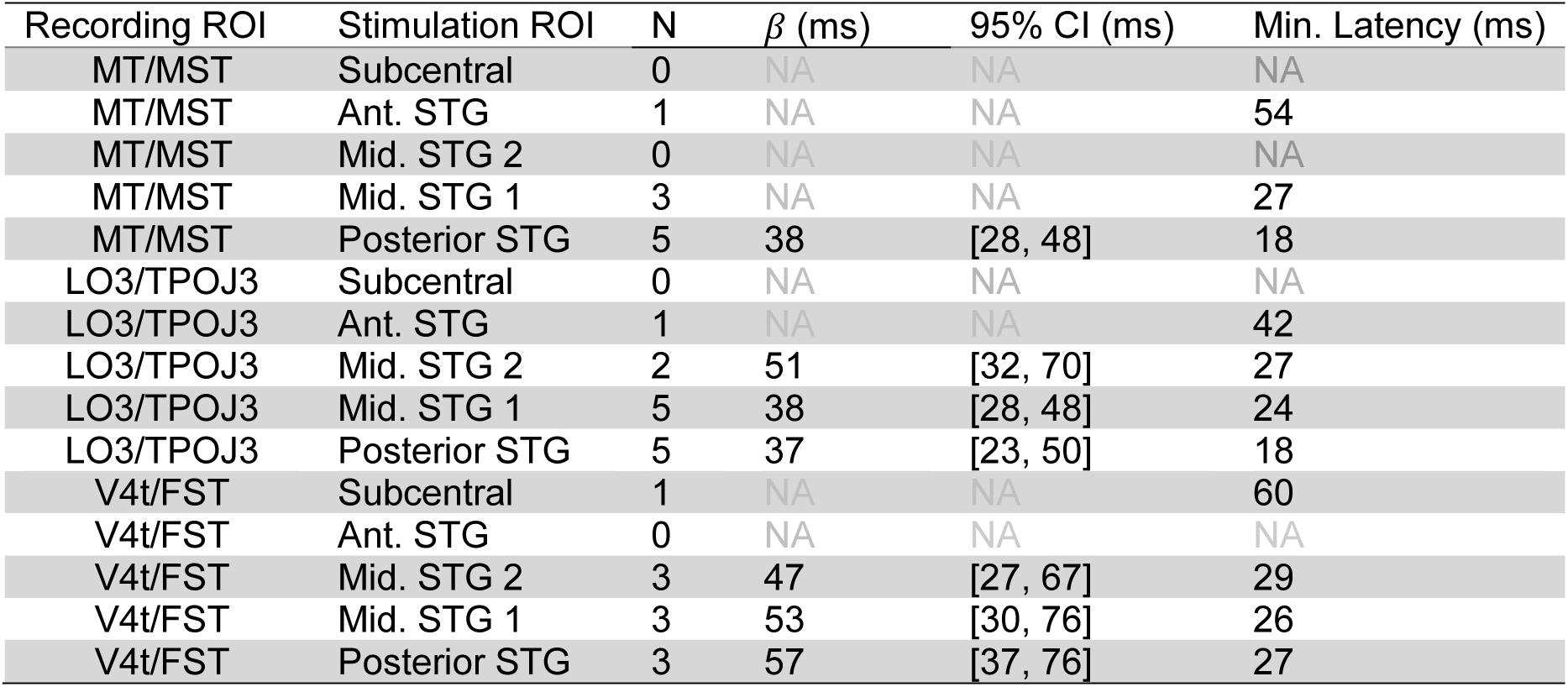
Latency estimates by stimulation and recording ‘subregion’ within V5/hMT+ ROI.

**Figure S1.**
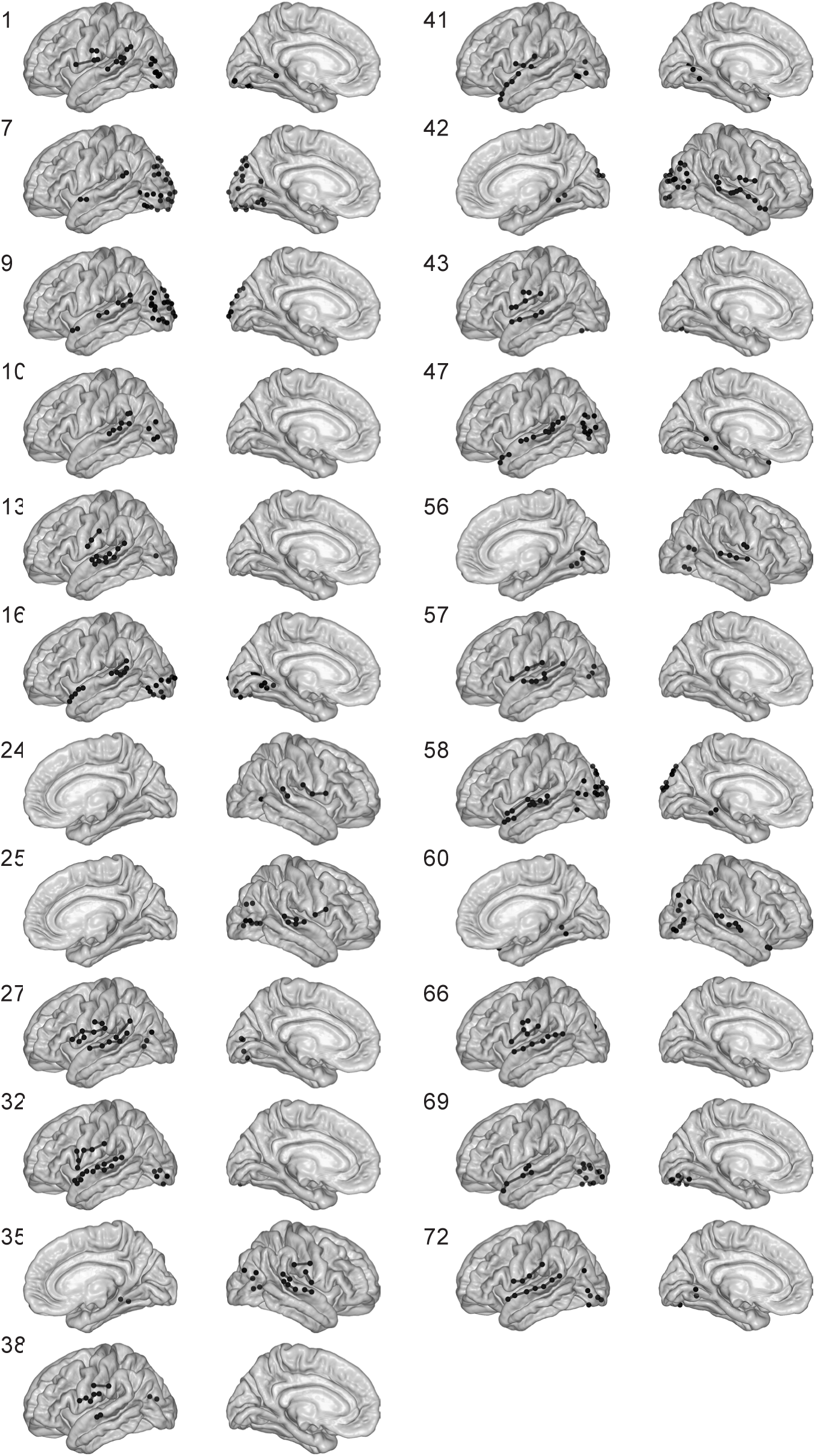
Included sites and stimulation pairs by patient. IDs correspond to index in original repository. Locations are displayed on the fsaverage brain. Lines link stimulation pairs. No patients had contralateral combinations of stimulation and recording site. Six patients had right-hemispheric coverage.

**Figure S2.**
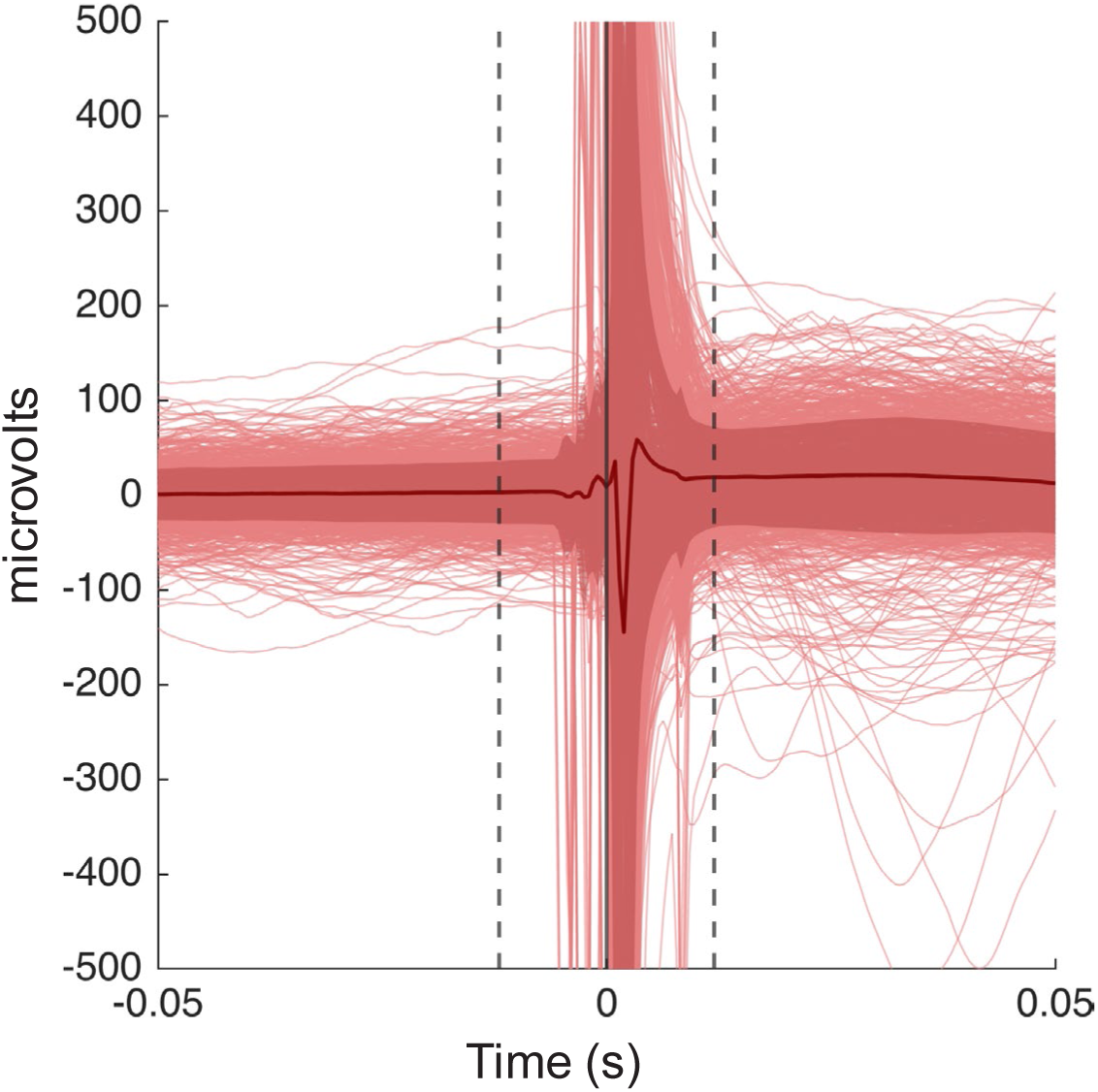
Averaged CCEP waveforms. All CCEP waveforms included in this study, and their averaged response (dark red line). Dashed lines highlight the range (-12 to 12ms) considered at risk of contamination by the stimulation artifact. Ribbon indicates standard deviation of the averaged response.

**Figure S3.**
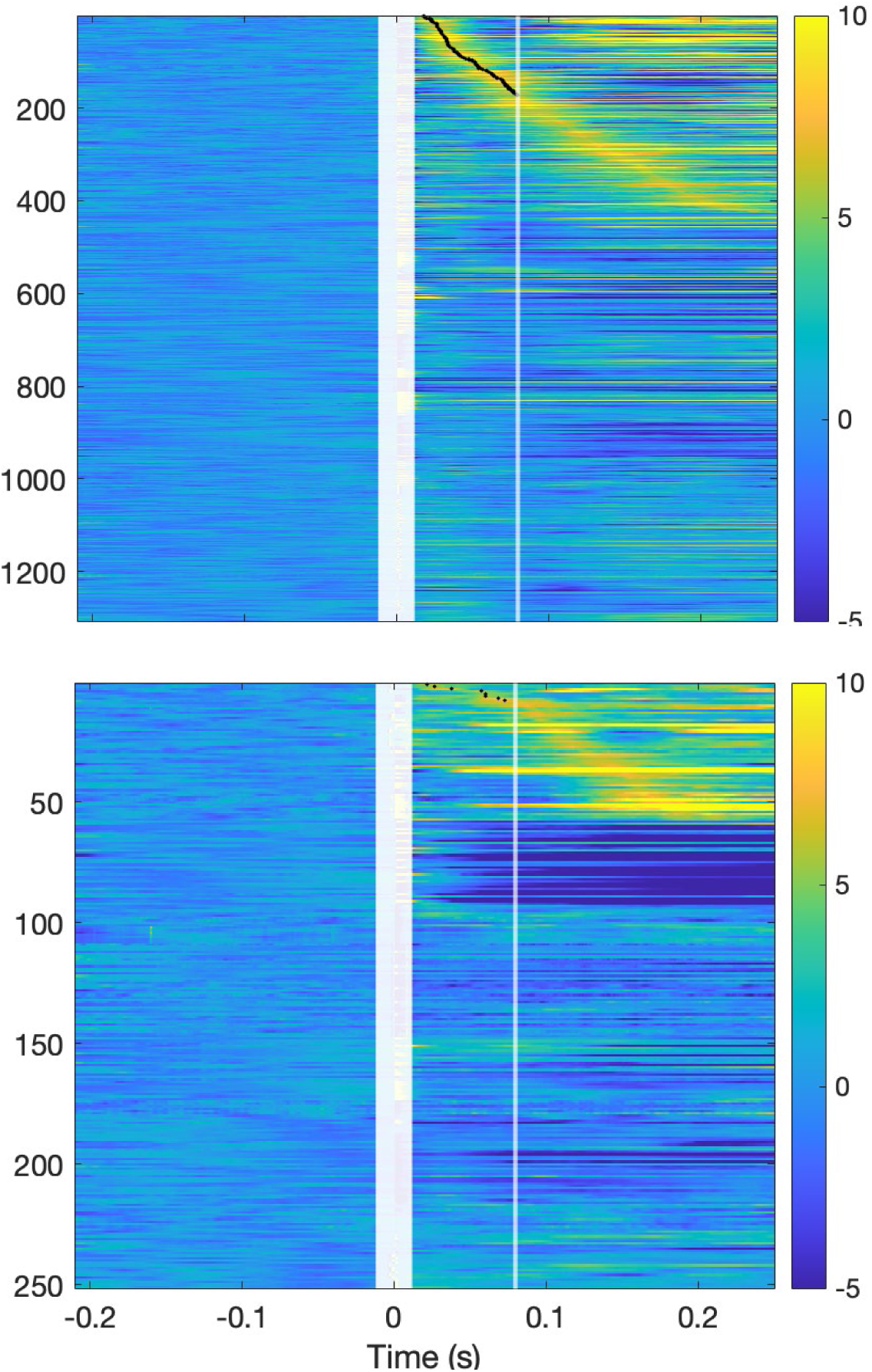
All CCEP responses. (Top) All occipital responses to STG stimulation, across all patients (z-scored waveforms), ordered by peak latency for detected responses. N1 responses are marked with a black dot. Vertical white lines indicate (a) the stimulation artifact window (fat line), and (b) the cutoff for N1 definition (80ms, thin line). **(Bottom)** All occipital responses to Subcentral stimulation.

**Figure S4.**
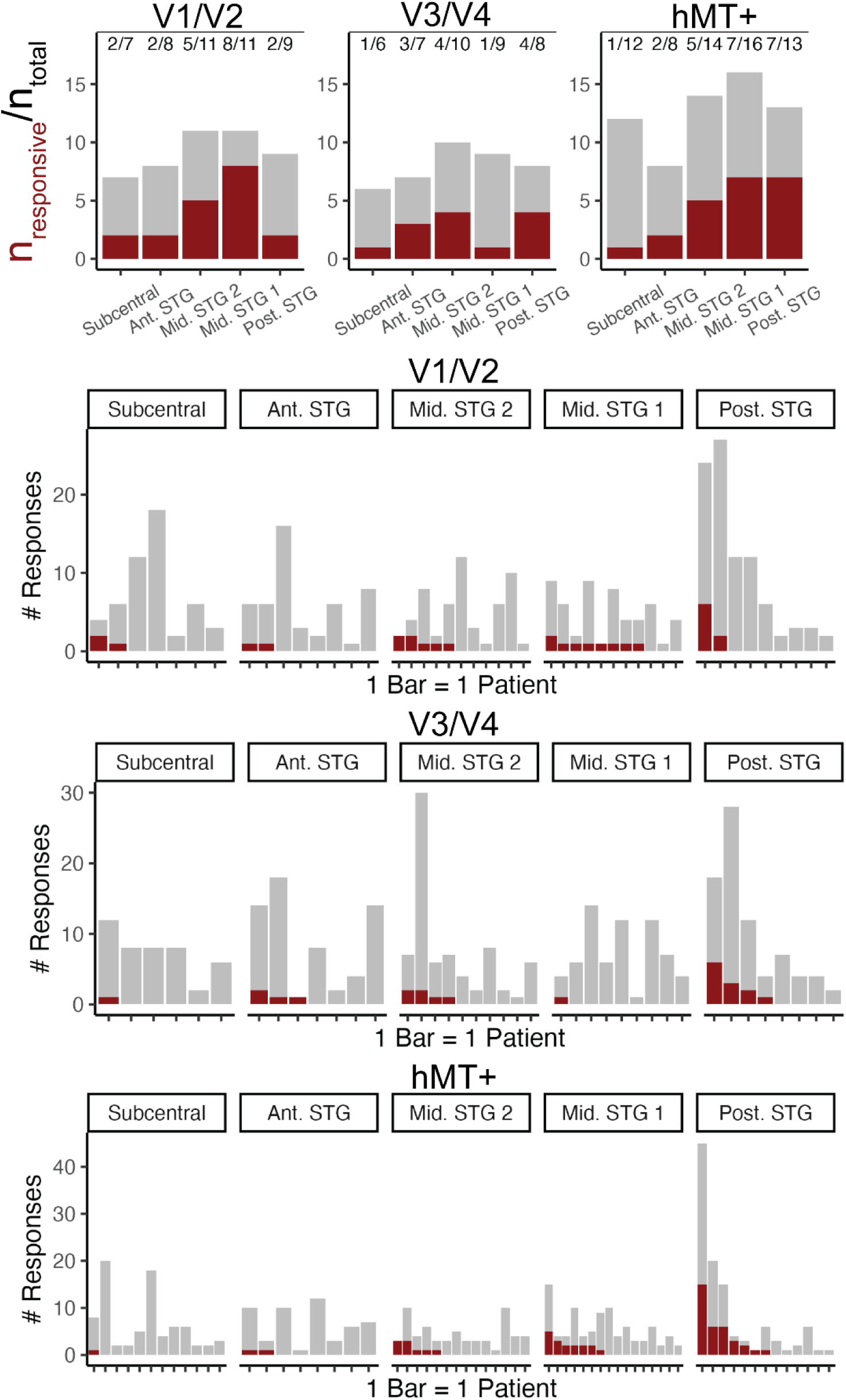
(Top row) Counts of patients for whom at least one response was detected (red) out of total number of patients (grey) for each region and stimulation site. (Remaining rows) Visualization of number of detected responses per patient (red) relative to total number of waveforms (grey). Each bar represents a single patient. Patients are sorted left-to-right based on the number of responses detected within each plot.

**Figure S5.**
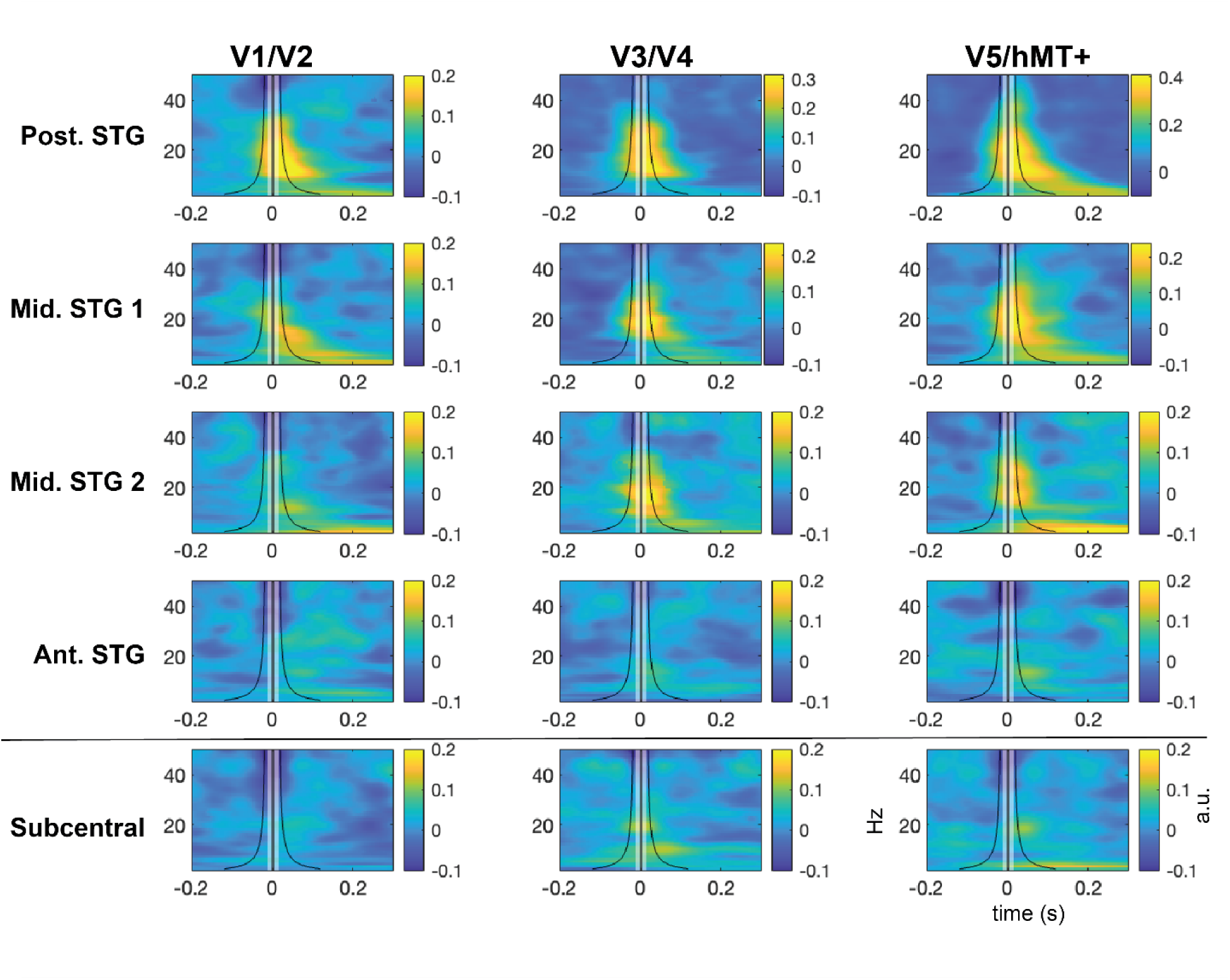
Time-frequency representation of all CCEPs from V1/V2 (left), V3/V4 (middle), and V5/hMT+ (right) ROIs. Rows indicate stimulation region. Heatmaps indicate power relative to prestimulus baseline for frequencies up to 50 Hz. The white bar indicates the region of interpolation. Black lines illustrate the standard deviation of the gaussian window used for wavelet generation at each frequency. Single-trial spectral information was modeled with a mixed-effects approach to generate fixed-effect estimates of power relative to baseline for each time/frequency point. For these models (with trial nested within electrode nested within subject), the random effects structure included crossed random effects of electrode, run (for patients with multiple runs), and stimulation pair.

**Figure S6.**
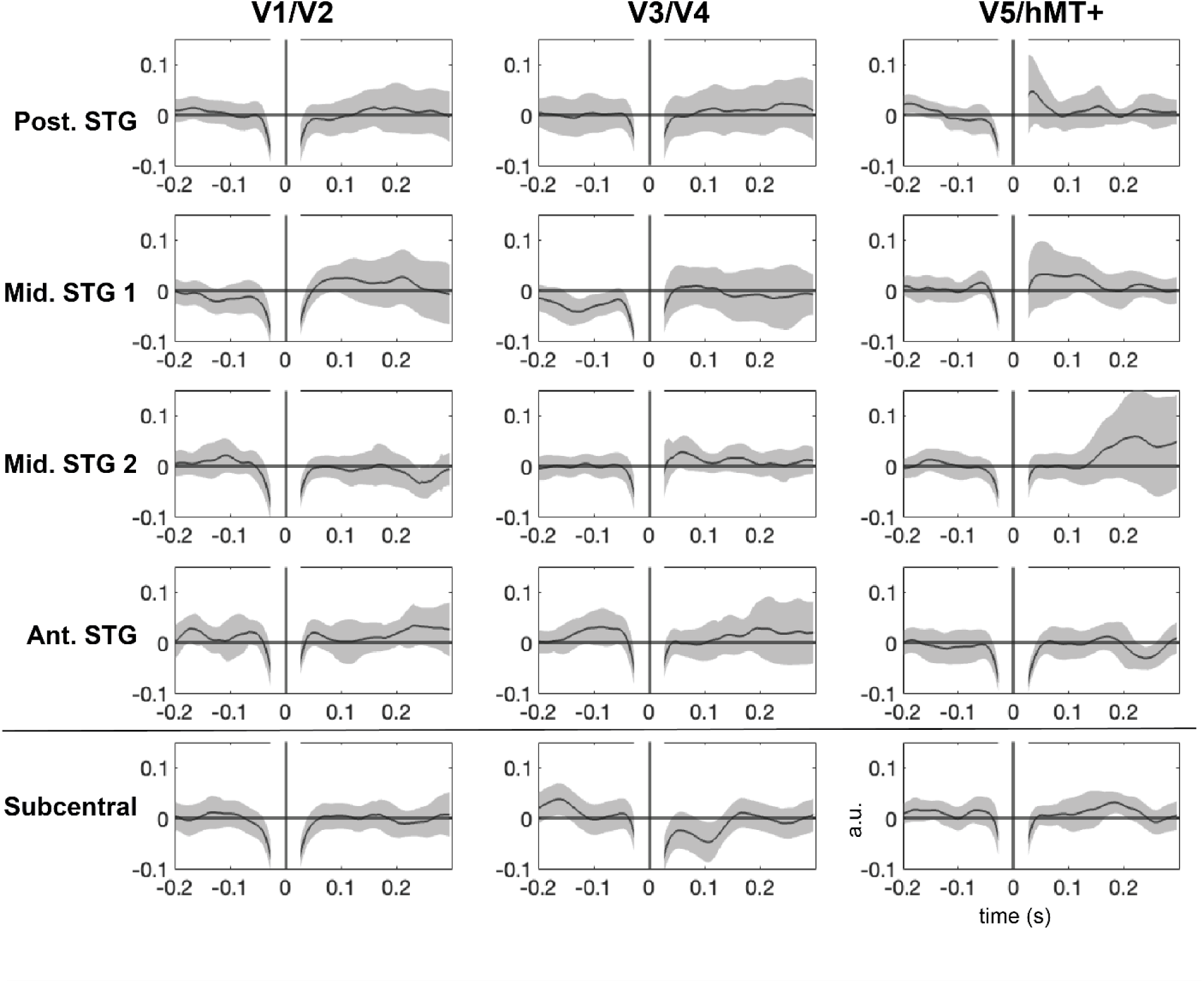
Time-series representation of high-gamma power for all CCEPs from V1/V2 (left), V3/V4 (middle), and V5/hMT+ (right) ROIs. Rows indicate stimulation region. Lines indicate high-gamma power relative to prestimulus baseline. The approximate range influenced by interpolation of the stimulation artifact is masked surrounding 0 (+/-1 standard deviation of the gaussian window used for wavelet generation). As in Figure S5, single-trial spectral information was modeled with a mixed-effects approach to generate fixed-effect estimates of power relative to baseline for each time/frequency point. The black line indicates the fixed intercept of the model at a given timepoint, and the ribbons indicate the 95% CI surrounding the fixed effect.

## References

Allman, B. L., & Meredith, M. A. (2007). Multisensory Processing in “Unimodal” Neurons: Cross-Modal Subthreshold Auditory Effects in Cat Extrastriate Visual Cortex. Journal of Neurophysiology, 98(1), 545–549. 10.1152/jn.00173.2007

Araki, K., Terada, K., Usui, K., Usui, N., Araki, Y., Baba, K., Matsuda, K., Tottori, T., & Inoue, Y. (2015). Bidirectional neural connectivity between basal temporal and posterior language areas in humans. Clinical Neurophysiology, 126(4), 682–688. 10.1016/j.clinph.2014.07.020

Bastos, A. M., Vezoli, J., Bosman, C. A., Schoffelen, J.-M., Oostenveld, R., Dowdall, J. R., De Weerd, P., Kennedy, H., & Fries, P. (2015). Visual Areas Exert Feedforward and Feedback Influences through Distinct Frequency Channels. Neuron, 85(2), 390–401. 10.1016/j.neuron.2014.12.018

Bates, D., Mächler, M., Bolker, B., & Walker, S. (2015). Fitting Linear Mixed-Effects Models Using lme4. Journal of Statistical Software, 67, 1–48. 10.18637/jss.v067.i01

Bedny, M., Konkle, T., Pelphrey, K., Saxe, R., & Pascual-Leone, A. (2010). Sensitive Period for a Multimodal Response in Human Visual Motion Area MT/MST. Current Biology, 20(21), 1900–1906. 10.1016/j.cub.2010.09.044

Beer, A. L., Plank, T., & Greenlee, M. W. (2011). Diffusion tensor imaging shows white matter tracts between human auditory and visual cortex. Experimental Brain Research, 213(2), 299–308. 10.1007/s00221-011-2715-y

Bieser, A., & Müller-Preuss, P. (1996). Auditory responsive cortex in the squirrel monkey: Neural responses to amplitude-modulated sounds. Experimental Brain Research, 108(2), 273–284. 10.1007/BF00228100

Borra, E., & Rockland, K. S. (2011). Projections to Early Visual Areas V1 and V2 in the Calcarine Fissure from Parietal Association Areas in the Macaque. Frontiers in Neuroanatomy, 5. 10.3389/fnana.2011.00035

Brang, D., Plass, J., Sherman, A., Stacey, W. C., Wasade, V. S., Grabowecky, M., Ahn, E., Towle, V. L., Tao, J. X., Wu, S., Issa, N. P., & Suzuki, S. (2022). Visual cortex responds to sound onset and offset during passive listening. Journal of Neurophysiology, 127(6), 1547–1563. 10.1152/jn.00164.2021

Brang, D., Towle, V. L., Suzuki, S., Hillyard, S. A., Di Tusa, S., Dai, Z., Tao, J., Wu, S., & Grabowecky, M. (2015). Peripheral sounds rapidly activate visual cortex: Evidence from electrocorticography. Journal of Neurophysiology, 114(5), 3023–3028. 10.1152/jn.00728.2015

Buffalo, E. A., Fries, P., Landman, R., Buschman, T. J., & Desimone, R. (2011). Laminar differences in gamma and alpha coherence in the ventral stream. Proceedings of the National Academy of Sciences, 108(27), 11262–11267. 10.1073/pnas.1011284108

Cappe, C., Rouiller, E. M., & Barone, P. (2009). Multisensory anatomical pathways. Hearing Research, 258(1), 28–36. 10.1016/j.heares.2009.04.017

Clavagnier, S., Falchier, A., & Kennedy, H. (2004). Long-distance feedback projections to area V1: Implications for multisensory integration, spatial awareness, and visual consciousness. *Cognitive, Affective*, & Behavioral Neuroscience, 4(2), 117–126. 10.3758/CABN.4.2.117

Cunningham, E. C., Wang, R. F., & Beck, D. M. (2022). Effects of rhythmic auditory stimulation on vision: Oscillations in performance can be enhanced, but not induced. Journal of Experimental Psychology: Human Perception and Performance, 48(11), 1153–1171. 10.1037/xhp0001029

Destrieux, C., Fischl, B., Dale, A., & Halgren, E. (2010). Automatic parcellation of human cortical gyri and sulci using standard anatomical nomenclature. NeuroImage, 53(1), 1–15. 10.1016/j.neuroimage.2010.06.010

Dormal, G., Rezk, M., Yakobov, E., Lepore, F., & Collignon, O. (2016). Auditory motion in the sighted and blind: Early visual deprivation triggers a large-scale imbalance between auditory and “visual” brain regions. NeuroImage, 134, 630–644. 10.1016/j.neuroimage.2016.04.027

Falchier, A., Cappe, C., Barone, P., & Schroeder, C. E. (2012). Sensory Convergence in Low-Level Cortices. In B. E. Stein (Ed.), The New Handbook of Multisensory Processing (pp. 67–80). The MIT Press. 10.7551/mitpress/8466.003.0008

Falchier, A., Clavagnier, S., Barone, P., & Kennedy, H. (2002). Anatomical Evidence of Multimodal Integration in Primate Striate Cortex. Journal of Neuroscience, 22(13), 5749– 5759. 10.1523/JNEUROSCI.22-13-05749.2002

Ferraro, S., Van Ackeren, M. J., Mai, R., Tassi, L., Cardinale, F., Nigri, A., Bruzzone, M. G., D’Incerti, L., Hartmann, T., Weisz, N., & Collignon, O. (2020). Stereotactic electroencephalography in humans reveals multisensory signal in early visual and auditory cortices. Cortex, 126, 253–264. 10.1016/j.cortex.2019.12.032

Frassinetti, F., Bolognini, N., & Làdavas, E. (2002). Enhancement of visual perception by crossmodal visuo-auditory interaction. Experimental Brain Research, 147(3), 332–343. 10.1007/s00221-002-1262-y

Giard, M. H., & Peronnet, F. (1999). Auditory-Visual Integration during Multimodal Object Recognition in Humans: A Behavioral and Electrophysiological Study. Journal of Cognitive Neuroscience, 11(5), 473–490. 10.1162/089892999563544

Glasser, M. F., Coalson, T. S., Robinson, E. C., Hacker, C. D., Harwell, J., Yacoub, E., Ugurbil, K., Andersson, J., Beckmann, C. F., Jenkinson, M., Smith, S. M., & Van Essen, D. C. (2016). A multi-modal parcellation of human cerebral cortex. Nature, 536(7615), 171– 178. 10.1038/nature18933

Gurtubay-Antolin, A., Battal, C., Maffei, C., Rezk, M., Mattioni, S., Jovicich, J., & Collignon, O. (2021). Direct Structural Connections between Auditory and Visual Motion-Selective Regions in Humans. Journal of Neuroscience, 41(11), 2393–2405. 10.1523/JNEUROSCI.1552-20.2021

Harms, M. P., & Melcher, J. R. (2002). Sound Repetition Rate in the Human Auditory Pathway: Representations in the Waveshape and Amplitude of fMRI Activation. Journal of Neurophysiology, 88(3), 1433–1450. 10.1152/jn.2002.88.3.1433

Huang, H., Gregg, N. M., Ojeda Valencia, G., Brinkmann, B. H., Lundstrom, B. N., Worrell, G. A., Miller, K. J., & Hermes, D. (2023). Electrical Stimulation of Temporal and Limbic Circuitry Produces Distinct Responses in Human Ventral Temporal Cortex. The Journal of Neuroscience, 43(24), 4434–4447. 10.1523/JNEUROSCI.1325-22.2023

Huang, T., Chen, X., Jiang, J., Zhen, Z., & Liu, J. (2019). A probabilistic atlas of the human motion complex built from large-scale functional localizer data. Human Brain Mapping, 40(12), 3475–3487. 10.1002/hbm.24610

Keller, C. J., Honey, C. J., Mégevand, P., Entz, L., Ulbert, I., & Mehta, A. D. (2014). Mapping human brain networks with cortico-cortical evoked potentials. Philosophical Transactions of the Royal Society B: Biological Sciences, 369(1653), 20130528. 10.1098/rstb.2013.0528

Lakatos, P., O’Connell, M. N., Barczak, A., Mills, A., Javitt, D. C., & Schroeder, C. E. (2009). The Leading Sense: Supramodal Control of Neurophysiological Context by Attention. Neuron, 64(3), 419–430. 10.1016/j.neuron.2009.10.014

Lemaréchal, J.-D., Jedynak, M., Trebaul, L., Boyer, A., Tadel, F., Bhattacharjee, M., Deman, P., Tuyisenge, V., Ayoubian, L., Hugues, E., Chanteloup-Forêt, B., Saubat, C., Zouglech, R., Reyes Mejia, G. C., Tourbier, S., Hagmann, P., Adam, C., Barba, C., Bartolomei, F., Nacci, E. (2022). A brain atlas of axonal and synaptic delays based on modelling of cortico-cortical evoked potentials. Brain, 145(5), 1653–1667. 10.1093/brain/awab362

Liégeois-Chauvel, C., Musolino, A., Badier, J. M., Marquis, P., & Chauvel, P. (1994). Evoked potentials recorded from the auditory cortex in man: Evaluation and topography of the middle latency components. Electroencephalography and Clinical Neurophysiology/Evoked Potentials Section, 92(3), 204–214. 10.1016/0168-5597(94)90064-7

Lyon, D. C., Nassi, J. J., & Callaway, E. M. (2010). A Disynaptic Relay from Superior Colliculus to Dorsal Stream Visual Cortex in Macaque Monkey. Neuron, 65(2), 270–279. 10.1016/j.neuron.2010.01.003

Majka, P., Rosa, M. G. P., Bai, S., Chan, J. M., Huo, B.-X., Jermakow, N., Lin, M. K., Takahashi, Y. S., Wolkowicz, I. H., Worthy, K. H., Rajan, R., Reser, D. H., Wójcik, D. K., Okano, H., & Mitra, P. P. (2019). Unidirectional monosynaptic connections from auditory areas to the primary visual cortex in the marmoset monkey. Brain Structure and Function, 224(1), 111–131. 10.1007/s00429-018-1764-4

Matsumoto, R., Kunieda, T., & Nair, D. (2017). Single pulse electrical stimulation to probe functional and pathological connectivity in epilepsy. Seizure, 44, 27–36. 10.1016/j.seizure.2016.11.003

Matsumoto, R., Nair, D. R., Ikeda, A., Fumuro, T., LaPresto, E., Mikuni, N., Bingaman, W., Miyamoto, S., Fukuyama, H., Takahashi, R., Najm, I., Shibasaki, H., & Lüders, H. O. (2012). Parieto-frontal network in humans studied by cortico-cortical evoked potential. Human Brain Mapping, 33(12), 2856–2872. 10.1002/hbm.21407

Matsumoto, R., Nair, D. R., LaPresto, E., Najm, I., Bingaman, W., Shibasaki, H., & Lüders, H. O. (2004). Functional connectivity in the human language system: A cortico-cortical evoked potential study. Brain, 127(10), 2316–2330. 10.1093/brain/awh246

McDonald, J. J., Teder-Sälejärvi, W. A., & Hillyard, S. A. (2000). Involuntary orienting to sound improves visual perception. Nature, 407(6806), 906–908. 10.1038/35038085

Mercier, M. R., Foxe, J. J., Fiebelkorn, I. C., Butler, J. S., Schwartz, T. H., & Molholm, S. (2013). Auditory-driven phase reset in visual cortex: Human electrocorticography reveals mechanisms of early multisensory integration. NeuroImage, 79, 19–29. 10.1016/j.neuroimage.2013.04.060

Meredith, M. A., Allman, B. L., Keniston, L. P., & Clemo, H. R. (2009). Auditory influences on non-auditory cortices. Hearing Research, 258(1), 64–71. 10.1016/j.heares.2009.03.005

Miller, K. J., Müller, K.-R., & Hermes, D. (2021). Basis profile curve identification to understand electrical stimulation effects in human brain networks. PLOS Computational Biology, 17(9), e1008710. 10.1371/journal.pcbi.1008710

Mishra, J., Martinez, A., Sejnowski, T. J., & Hillyard, S. A. (2007). Early Cross-Modal Interactions in Auditory and Visual Cortex Underlie a Sound-Induced Visual Illusion. Journal of Neuroscience, 27(15), 4120–4131. 10.1523/JNEUROSCI.4912-06.2007

Molholm, S., Ritter, W., Murray, M. M., Javitt, D. C., Schroeder, C. E., & Foxe, J. J. (2002). Multisensory auditory–visual interactions during early sensory processing in humans: A high-density electrical mapping study. Cognitive Brain Research, 14(1), 115–128. 10.1016/S0926-6410(02)00066-6

Naue, N., Rach, S., Struber, D., Huster, R. J., Zaehle, T., Korner, U., & Herrmann, C. S. (2011). Auditory Event-Related Response in Visual Cortex Modulates Subsequent Visual Responses in Humans. Journal of Neuroscience, 31(21), 7729–7736. 10.1523/JNEUROSCI.1076-11.2011

Noesselt, T., Tyll, S., Boehler, C. N., Budinger, E., Heinze, H.-J., & Driver, J. (2010). Sound-Induced Enhancement of Low-Intensity Vision: Multisensory Influences on Human Sensory-Specific Cortices and Thalamic Bodies Relate to Perceptual Enhancement of Visual Detection Sensitivity. Journal of Neuroscience, 30(41), 13609–13623. 10.1523/JNEUROSCI.4524-09.2010

Palmer, S. M., & Rosa, M. G. P. (2006). Quantitative Analysis of the Corticocortical Projections to the Middle Temporal Area in the Marmoset Monkey: Evolutionary and Functional Implications. Cerebral Cortex, 16(9), 1361–1375. 10.1093/cercor/bhj078

Plass, J., Ahn, E., Towle, V. L., Stacey, W. C., Wasade, V. S., Tao, J., Wu, S., Issa, N. P., & Brang, D. (2019). Joint encoding of auditory timing and location in visual cortex. Journal of Cognitive Neuroscience, 31(7), 1002–1017. 10.1162/jocn_a_01399

Poirier, C., Collignon, O., DeVolder, A. G., Renier, L., Vanlierde, A., Tranduy, D., & Scheiber, C. (2005). Specific activation of the V5 brain area by auditory motion processing: An fMRI study. Cognitive Brain Research, 25(3), 650–658. 10.1016/j.cogbrainres.2005.08.015

Poirier, C., Collignon, O., Scheiber, C., Renier, L., Vanlierde, A., Tranduy, D., Veraart, C., & De Volder, A. G. (2006). Auditory motion perception activates visual motion areas in early blind subjects. NeuroImage, 31(1), 279–285. 10.1016/j.neuroimage.2005.11.036

Raij, T., Ahveninen, J., Lin, F.-H., Witzel, T., Jääskeläinen, I. P., Letham, B., Israeli, E., Sahyoun, C., Vasios, C., Stufflebeam, S., Hämäläinen, M., & Belliveau, J. W. (2010). Onset timing of cross-sensory activations and multisensory interactions in auditory and visual sensory cortices. European Journal of Neuroscience, 31(10), 1772–1782. 10.1111/j.1460-9568.2010.07213.x

Rezk, M., Cattoir, S., Battal, C., Occelli, V., Mattioni, S., & Collignon, O. (2020). Shared Representation of Visual and Auditory Motion Directions in the Human Middle-Temporal Cortex. Current Biology, 30(12), 2289–2299.e8. 10.1016/j.cub.2020.04.039

Rockland, K. S., & Ojima, H. (2003). Multisensory convergence in calcarine visual areas in macaque monkey. International Journal of Psychophysiology, 50(1), 19–26. 10.1016/S0167-8760(03)00121-1

Romei, V., Gross, J., & Thut, G. (2012). Sounds reset rhythms of visual cortex and corresponding human visual perception. Current Biology: CB, 22(9), 807–813. 10.1016/j.cub.2012.03.025

Romei, V., Murray, M. M., Cappe, C., & Thut, G. (2009). Preperceptual and Stimulus-Selective Enhancement of Low-Level Human Visual Cortex Excitability by Sounds. Current Biology, 19(21), 1799–1805. 10.1016/j.cub.2009.09.027

Romei, V., Murray, M. M., Merabet, L. B., & Thut, G. (2007). Occipital Transcranial Magnetic Stimulation Has Opposing Effects on Visual and Auditory Stimulus Detection: Implications for Multisensory Interactions. Journal of Neuroscience, 27(43), 11465– 11472. 10.1523/JNEUROSCI.2827-07.2007

Saenz, M., Lewis, L. B., Huth, A. G., Fine, I., & Koch, C. (2008). Visual Motion Area MT+/V5 Responds to Auditory Motion in Human Sight-Recovery Subjects. Journal of Neuroscience, 28(20), 5141–5148. 10.1523/JNEUROSCI.0803-08.2008

Shams, L., Kamitani, Y., & Shimojo, S. (2000). What you see is what you hear. Nature, 408(6814), 788–788. 10.1038/35048669

Shipley, T. (1964). Auditory Flutter-Driving of Visual Flicker. Science, 145(3638), 1328–1330. 10.1126/science.145.3638.1328

Simonyan, K., & Jürgens, U. (2002). Cortico-cortical projections of the motorcortical larynx area in the rhesus monkey. Brain Research, 949(1), 23–31. 10.1016/S0006-8993(02)02960-8

Simonyan, K., & Jürgens, U. (2003). Efferent subcortical projections of the laryngeal motorcortex in the rhesus monkey. Brain Research, 974(1), 43–59. 10.1016/S0006-8993(03)02548-4

Strnad, L., Peelen, M. V., Bedny, M., & Caramazza, A. (2013). Multivoxel Pattern Analysis Reveals Auditory Motion Information in MT+ of Both Congenitally Blind and Sighted Individuals. PLOS ONE, 8(4), e63198. 10.1371/journal.pone.0063198

Thorne, J. D., & Debener, S. (2014). Look now and hear what’s coming: On the functional role of cross-modal phase reset. Hearing Research, 307, 144–152. 10.1016/j.heares.2013.07.002

Trebaul, L., Deman, P., Tuyisenge, V., Jedynak, M., Hugues, E., Rudrauf, D., Bhattacharjee, M., Tadel, F., Chanteloup-Foret, B., Saubat, C., Reyes Mejia, G. C., Adam, C., Nica, A., Pail, M., Dubeau, F., Rheims, S., Trébuchon, A., Wang, H., Liu, S., … David, O. (2018). Probabilistic functional tractography of the human cortex revisited. NeuroImage, 181, 414–429. 10.1016/j.neuroimage.2018.07.039

van Blooijs, D., van den Boom, M. A., van der Aar, J. F., Huiskamp, G. M., Castegnaro, G., Demuru, M., Zweiphenning, W. J. E. M., van Eijsden, P., Miller, K. J., Leijten, F. S. S., & Hermes, D. (2023). Developmental trajectory of transmission speed in the human brain. Nature Neuroscience, 26(4), Article 4. 10.1038/s41593-023-01272-0

Vetter, P., Bola, Ł., Reich, L., Bennett, M., Muckli, L., & Amedi, A. (2020). Decoding Natural Sounds in Early “Visual” Cortex of Congenitally Blind Individuals. Current Biology, 30(15), 3039–3044.e2. 10.1016/j.cub.2020.05.071

Vroomen, J., & Gelder, B. de. (2000). Sound enhances visual perception: Cross-modal effects of auditory organization on vision. Journal of Experimental Psychology: Human Perception and Performance, 26(5), 1583–1590. 10.1037/0096-523.26.5.1583

Wallace, M. T., Wilkinson, L. K., & Stein, B. E. (1996). Representation and integration of multiple sensory inputs in primate superior colliculus. Journal of Neurophysiology. 10.1152/jn.1996.76.2.1246

Wang, Y., Celebrini, S., Trotter, Y., & Barone, P. (2008). Visuo-auditory interactions in the primary visual cortex of the behaving monkey: Electrophysiological evidence. BMC Neuroscience, 9(1), 79. 10.1186/1471-2202-9-79

Yamao, Y., Matsumoto, R., Kunieda, T., Arakawa, Y., Kobayashi, K., Usami, K., Shibata, S., Kikuchi, T., Sawamoto, N., Mikuni, N., Ikeda, A., Fukuyama, H., & Miyamoto, S. (2014). Intraoperative dorsal language network mapping by using single-pulse electrical stimulation. Human Brain Mapping, 35(9), 4345–4361. 10.1002/hbm.22479

