## Supplemental Information for "Electrical stimulation of the superior temporal gyrus evokes rapid responses in human visual cortex"

**Table S1. Patient details**

| ID* | Age (y) | Gender | Hemi. | Sessions (n) | Sample rate (Hz) | Stim. Pairs**  (n_STG_ / n_subc._) | Recording sites (n) |
| --- | --- | --- | --- | --- | --- | --- | --- |
| 1 | 15 | male | L | 6 | 512/2048 | 4 / 3 | 13 |
| 7 | 10 | female | L | 2 | 2048 | 2 / 0 | 34 |
| 9 | 21 | male | L | 2 | 2048 | 4 / 0 | 21 |
| 10 | 15 | male | L | 2 | 2048 | 4 / 0 | 4 |
| 13 | 45 | male | L | 1 | 2048 | 8 / 2 | 1 |
| 16 | 42 | female | L | 1 | 2048 | 6 / 0 | 18 |
| 24 | 17 | male | R | 2 | 2048 | 1 / 2 | 1 |
| 25 | 16 | male | R | 1 | 2048 | 3 / 1 | 8 |
| 27 | 49 | male | L | 1 | 2048 | 5 / 6 | 8 |
| 32 | 8 | female | L | 2 | 2048 | 8 / 4 | 5 |
| 35 | 22 | female | R | 1 | 2048 | 4 / 2 | 7 |
| 38 | 21 | male | L | 1 | 2048 | 1 / 4 | 2 |
| 41 | 34 | male | L | 1 | 2048 | 5 / 2 | 7 |
| 42 | 17 | male | R | 1 | 2048 | 7 / 2 | 18 |
| 43 | 18 | female | L | 1 | 2048 | 3 / 4 | 2 |
| 47 | 30 | female | L | 1 | 2048 | 8 / 0 | 12 |
| 56 | 30 | female | R | 1 | 2048 | 3 / 1 | 8 |
| 57 | 30 | female | L | 1 | 2048 | 4 / 2 | 3 |
| 58 | 15 | female | L | 1 | 2048 | 10 / 0 | 20 |
| 60 | 15 | male | R | 1 | 2048 | 4 / 0 | 11 |
| 66 | 35 | male | L | 2 | 2048 | 5 / 4 | 1 |
| 69 | 51 | female | L | 1 | 2048 | 4 / 0 | 19 |
| 72 | 27 | female | L | 1 | 2048 | 6 / 3 | 8 |

*ID in original repository; **Counts of included pairs for STG and Subcentral stimulation sites

**Table S2. Latency estimates by stimulation and recording ‘subregion’ within V5/hMT+ ROI**

| Recording ROI | Stimulation ROI | N | 𝛽 (ms) | 95% CI (ms) | Min. Latency (ms) |
| --- | --- | --- | --- | --- | --- |
| MT/MST | Subcentral | 0 | NA | NA | NA |
| MT/MST | Ant. STG | 1 | NA | NA | 54 |
| MT/MST | Mid. STG 2 | 0 | NA | NA | NA |
| MT/MST | Mid. STG 1 | 3 | NA | NA | 27 |
| MT/MST | Posterior STG | 5 | 38 | [28, 48] | 18 |
| LO3/TPOJ3 | Subcentral | 0 | NA | NA | NA |
| LO3/TPOJ3 | Ant. STG | 1 | NA | NA | 42 |
| LO3/TPOJ3 | Mid. STG 2 | 2 | 51 | [32, 70] | 27 |
| LO3/TPOJ3 | Mid. STG 1 | 5 | 38 | [28, 48] | 24 |
| LO3/TPOJ3 | Posterior STG | 5 | 37 | [23, 50] | 18 |
| V4t/FST | Subcentral | 1 | NA | NA | 60 |
| V4t/FST | Ant. STG | 0 | NA | NA | NA |
| V4t/FST | Mid. STG 2 | 3 | 47 | [27, 67] | 29 |
| V4t/FST | Mid. STG 1 | 3 | 53 | [30, 76] | 26 |
| V4t/FST | Posterior STG | 3 | 57 | [37, 76] | 27 |

**Figure S1. Included sites and stimulation pairs by patient.** IDs correspond to index in original repository. Locations are displayed on the fsaverage brain. Lines link stimulation pairs. No patients had contralateral combinations of stimulation and recording site. Six patients had right-hemispheric coverage.

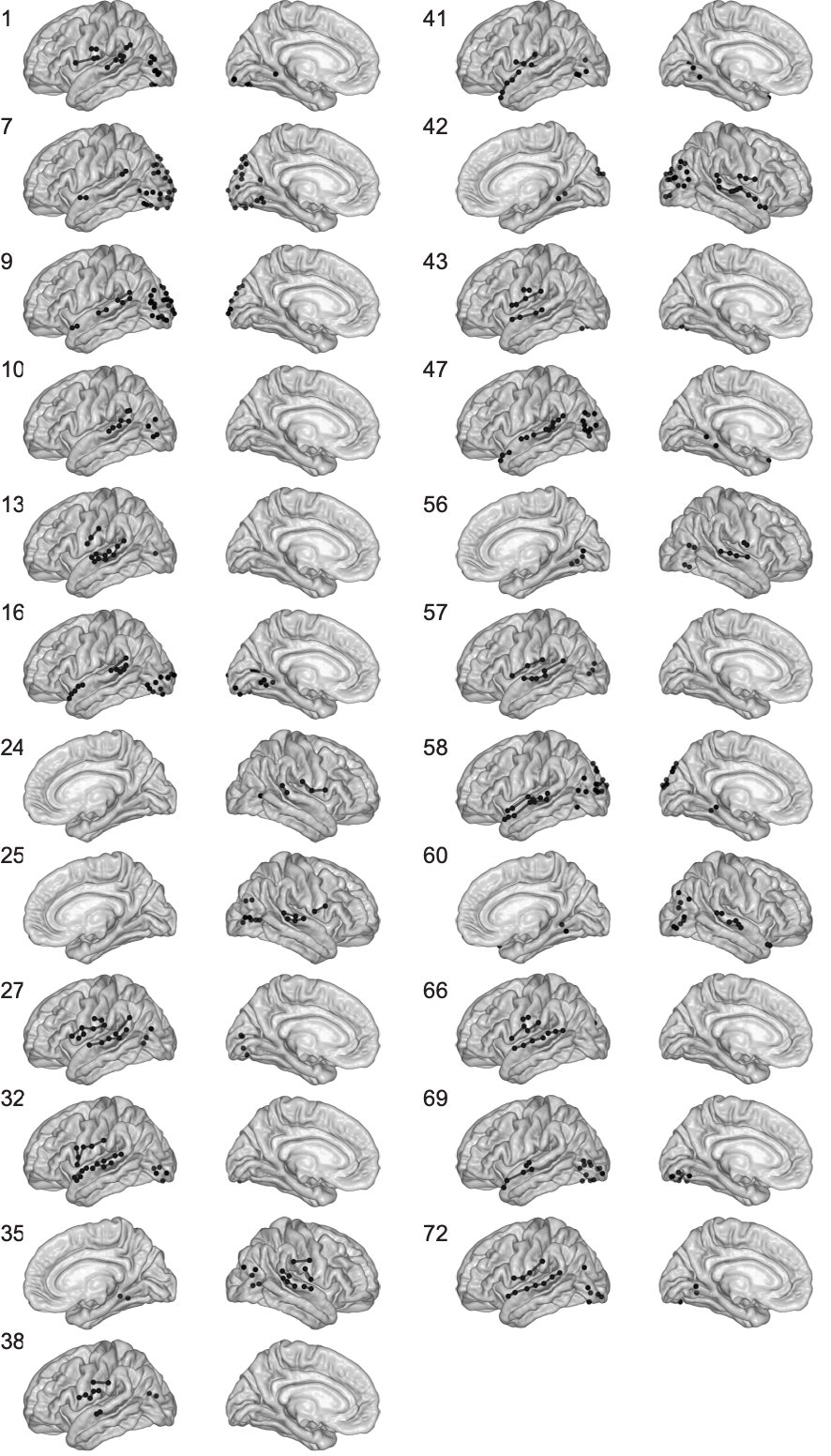

**Figure S2. Averaged CCEP waveforms.** All CCEP waveforms included in this study, and their averaged response (dark red line). Dashed lines highlight the range (-12 to 12ms) considered at risk of contamination by the stimulation artifact. Ribbon indicates standard deviation of the averaged response.

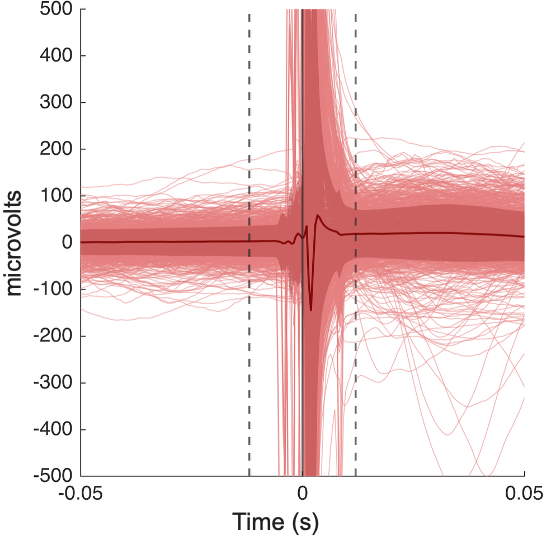

**Figure S3. All CCEP responses. (Top)** All occipital responses to STG stimulation, across all patients (z-scored waveforms), ordered by peak latency for detected responses. N1 responses are marked with a black dot. Vertical white lines indicate (a) the stimulation artifact window (fat line), and (b) the cutoff for N1 definition (80ms, thin line). **(Bottom)** All occipital responses to Subcentral stimulation.

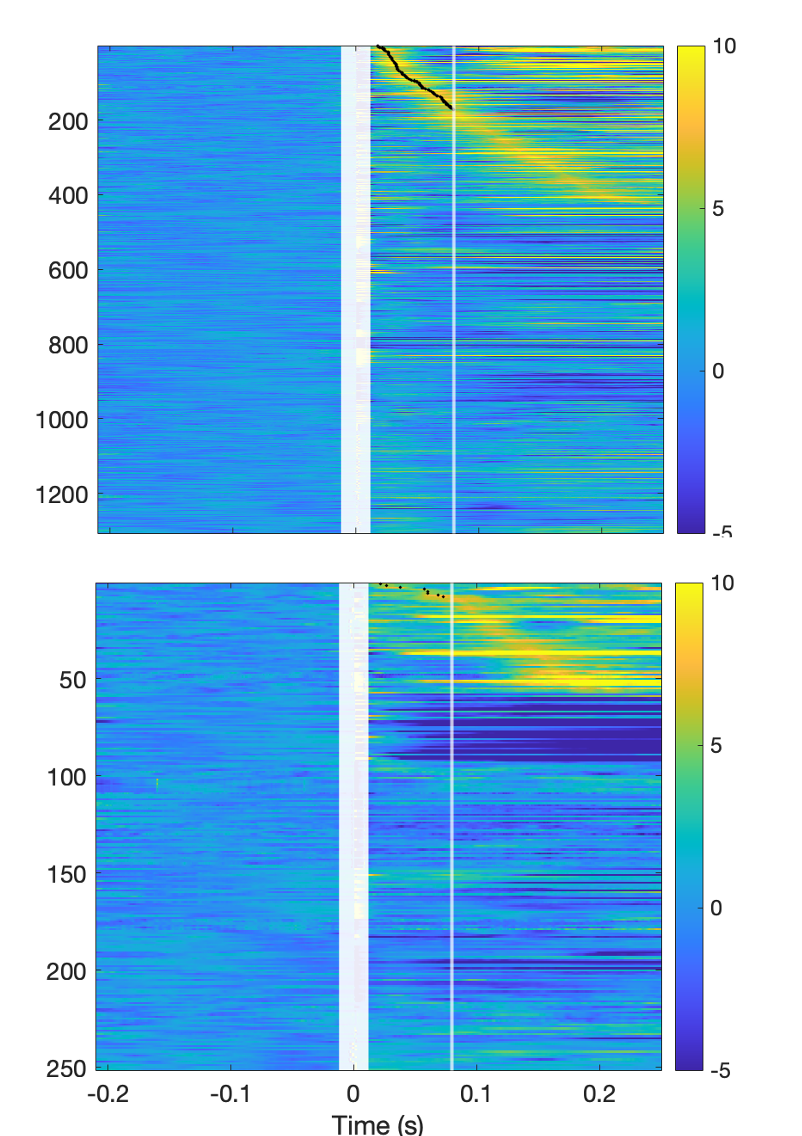

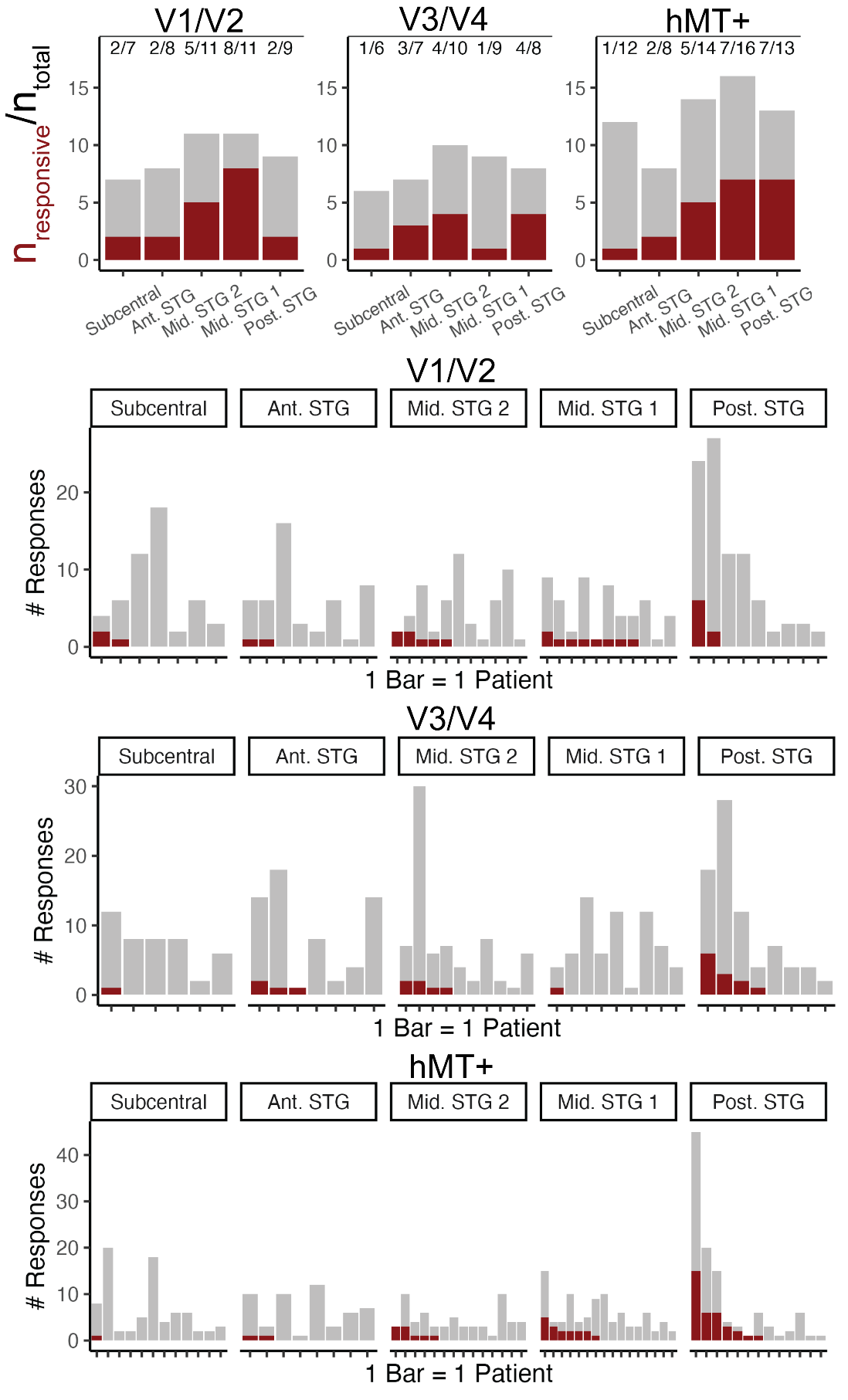
***Figure S4.*** *(Top row)* *Counts of patients for whom at least one response was detected (red) out of total number of patients (grey) for each region and stimulation site. (Remaining rows) Visualization of number of detected responses per patient (red) relative to total number of waveforms (grey). Each bar represents a single patient. Patients are sorted left-to-right based on the number of responses detected within each plot.*

***Figure S5.*** *Time-frequency representation of all CCEPs from V1/V2 (left), V3/V4 (middle), and V5/hMT+ (right) ROIs. Rows indicate stimulation region. Heatmaps indicate power relative to prestimulus baseline for frequencies up to 50 Hz. The white bar indicates the region of interpolation. Black lines illustrate the standard deviation of the gaussian window used for wavelet generation at each frequency. Single-trial spectral information was modeled with a mixed-effects approach to generate fixed-effect estimates of power relative to baseline for each time/frequency point. For these models (with trial nested within electrode nested within subject), the random effects structure included crossed random effects of electrode, run (for patients with multiple runs), and stimulation pair.*

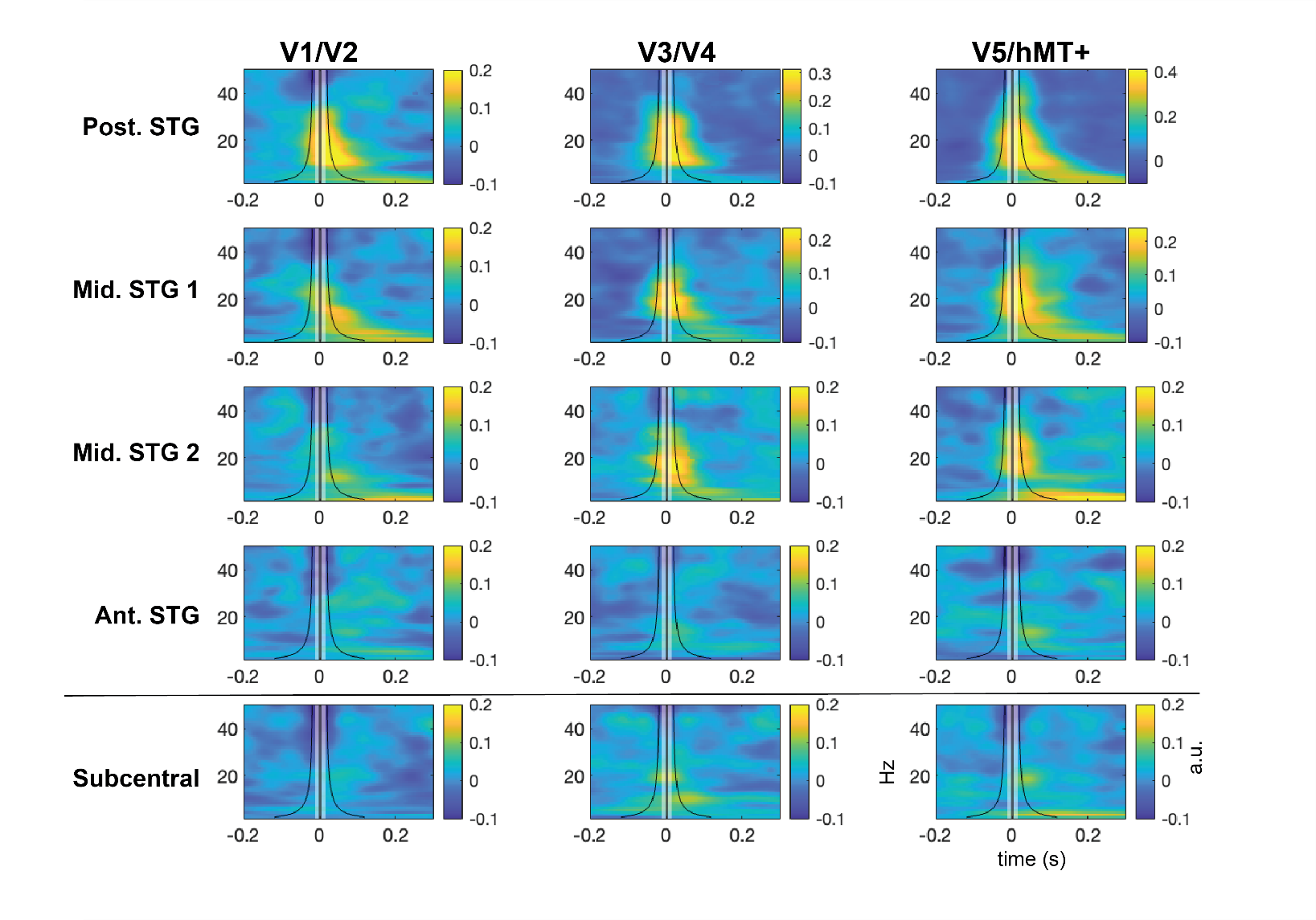

***Figure S6.*** *Time-series representation of high-gamma power for all CCEPs from V1/V2 (left), V3/V4 (middle), and V5/hMT+ (right) ROIs. Rows indicate stimulation region. Lines indicate high-gamma power relative to prestimulus baseline. The approximate range influenced by interpolation of the stimulation artifact is masked surrounding 0 (+/-1 standard deviation of the gaussian window used for wavelet generation). As in Figure S5, single-trial spectral information was modeled with a mixed-effects approach to generate fixed-effect estimates of power relative to baseline for each time/frequency point. The black line indicates the fixed intercept of the model at a given timepoint, and the ribbons indicate the 95% CI surrounding the fixed effect.*

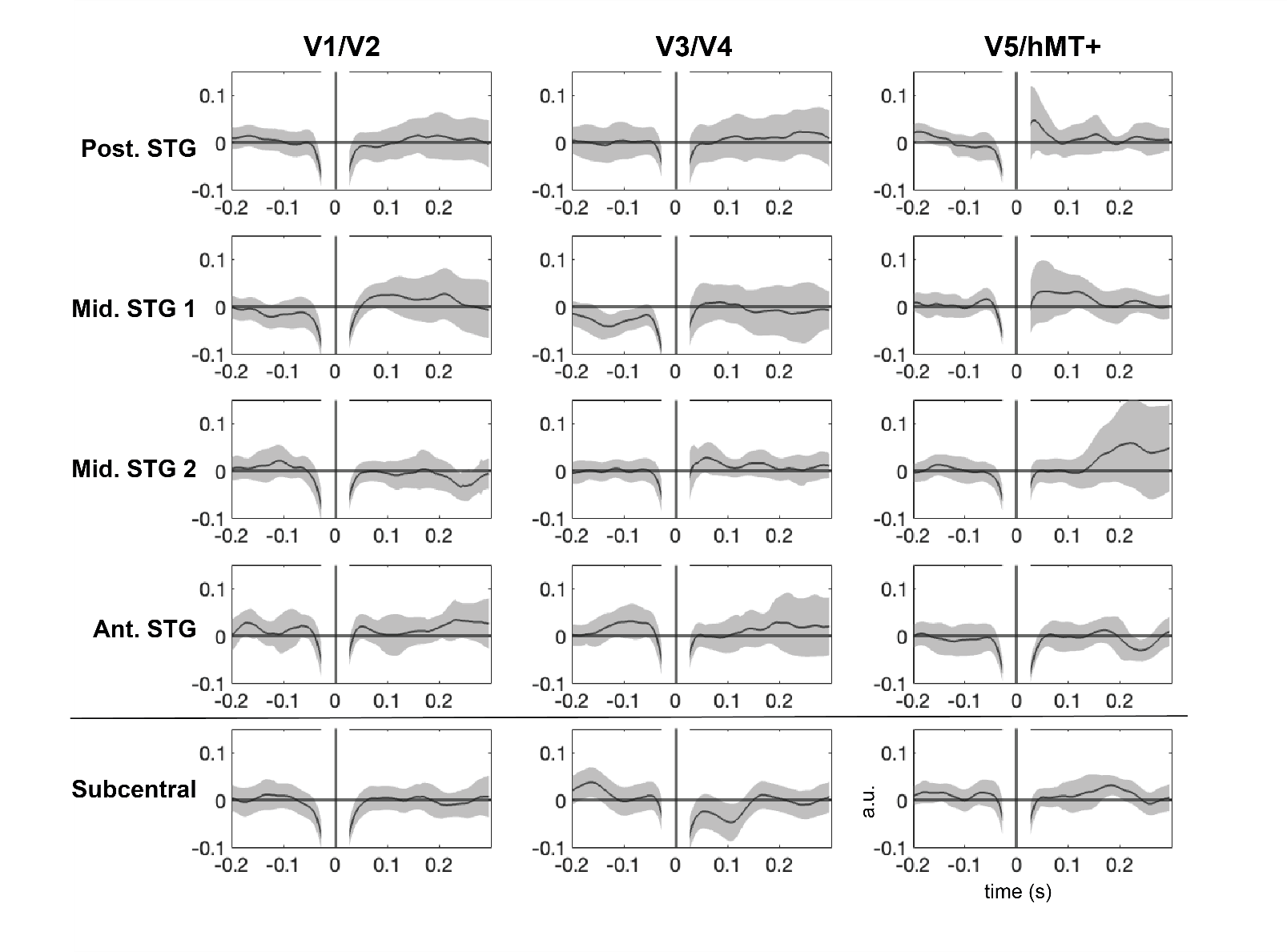
